# Live-Cell Chemoproteomic Profiling Identifies the Uncharacterised Protein YbaA as a Direct Target of Ciprofloxacin in *Escherichia coli*

**DOI:** 10.1101/2025.10.16.682360

**Authors:** Tricia C. Y. Seow, Jacob D. Bradbury, Emilia K. Taylor, Ludwig G. Bauer, Madia Harvey, Hildur Hreidarsdottir, Soraya M. Saghbini, Yiqiao Wang, Adam M. Thomas, Patricia Santos Barbosa, Kilian V. M. Huber, Georgia L. Isom, Thomas Lanyon-Hogg

## Abstract

Fluoroquinolone antibiotics, such as ciprofloxacin, are important broad-spectrum agents for a range of bacterial infections; however, fluoroquinolone usage is increasingly challenged by the emergence of resistance. Ciprofloxacin resistance mechanisms include mutations in the antibiotic target DNA gyrase, downregulation of porins required for bacterial cell penetration, and upregulation of efflux pumps to expel the antibiotic. However, new pathways driving bacterial tolerance to fluoroquinolones are still being discovered, suggesting that additional ciprofloxacin-binding proteins may exist in bacteria. In this study, we report the use of affinity-based protein profiling (AfBPP) with photo-crosslinking chemical probes to identify protein binding partners of ciprofloxacin in live *E. coli* cells. AfBPP identified novel ciprofloxacin binding proteins including YjdN and YbaA, whose molecular functions are as yet unannotated. Target engagement was validated using genetic knockout and biophysical binding assays, and key interactions identified in the ciprofloxacin binding site of YbaA. Collectively, this study demonstrates that additional and previously unreported biological interactions can exist for well-established antibiotics, and provides methodology to identify and interrogate these interactions in detail.

## Introduction

Antimicrobial resistance (AMR) is one of the most serious threats to health globally. In 2019, there were an estimated 1.27 million deaths directly attributable to bacterial AMR, a larger death toll than HIV/AIDS and malaria combined.^[1]^ Antibiotics are also essential for many frontline medical treatments, such as surgery and cancer chemotherapy, with AMR associated deaths estimated at 4.95 million in 2019 alone.^[1]^ The dissemination of AMR is accelerated by the misuse and overuse of antibiotics in both human healthcare and agriculture.^[2]^ Despite the pressing need for new antibiotics, the number of novel compounds currently in development is severely limited, as rapid resistance emergence means new antibiotics require safeguarding as ‘agents of last resort’, thus restricting their commercial market and disincentivising development.^[3]^ Given the increasingly challenging climate for development of new antibiotics with novel targets, it is increasingly important that current antibiotic mechanisms of action (MoA) and resistance are fully understood.

Ciprofloxacin (**CFX, 1**, Fig 1A) is a broad-spectrum fluoroquinolone antibiotic included in the World Health Organisation’s list of Essential Medicines,^[4]^ which is used to treat a range of infections including challenging Gram-negative pathogens such as *Escherichia coli* (*E. coli*).^[5]^ **CFX** inhibits two type IIA topoisomerases in bacteria to induce DNA double-strand breaks: DNA gyrase (GyrAB) is the primary target of **CFX** in *E. coli*, with topoisomerase IV (ParCE) a secondary target.^[6-7]^ Various **CFX** resistance mechanisms have been identified in *E. coli*, the most potent being the *gyrA* S83L mutation, which weakens the target interaction with **CFX**.^[8]^ Additional resistance mechanisms include decreased expression of OmpC and OmpF porin proteins proposed to allow **CFX** uptake into *E. coli*, or increased expression of efflux pumps such as AcrAB-TolC which reduce the intracellular **CFX** concentration.^[9]^ Formation of biofilms by bacteria can also increase antibiotic tolerance, and sub-minimum inhibitory concentrations (MICs) of **CFX** can promote or inhibit biofilm formation in different *E. coli* strains.^[10-12]^ Repair of **CFX**-induced DNA double strand breaks by RecBCD in *E. coli* activates the ‘SOS response’, triggering expression of error-prone polymerases, increasing horizontal gene transfer, and accelerating resistance.^[13-14]^ Inhibitors of **CFX** efflux or the SOS response in *Staphylococcus aureus* can resensitize pathogens to **CFX** and slow **CFX** resistance emergence,^[15-17]^ demonstrating the potential of antibiotic adjuvants to recover **CFX** activity. However, known resistance mechanisms only account for 50-70% of fluoroquinolone-resistant *E. coli* isolates^[18]^ and new pathways driving bacterial survival in the presence of quinolones are still being discovered,^[19]^ suggesting additional binding partners of **CFX** may remain to be discovered.

**Figure 1.**
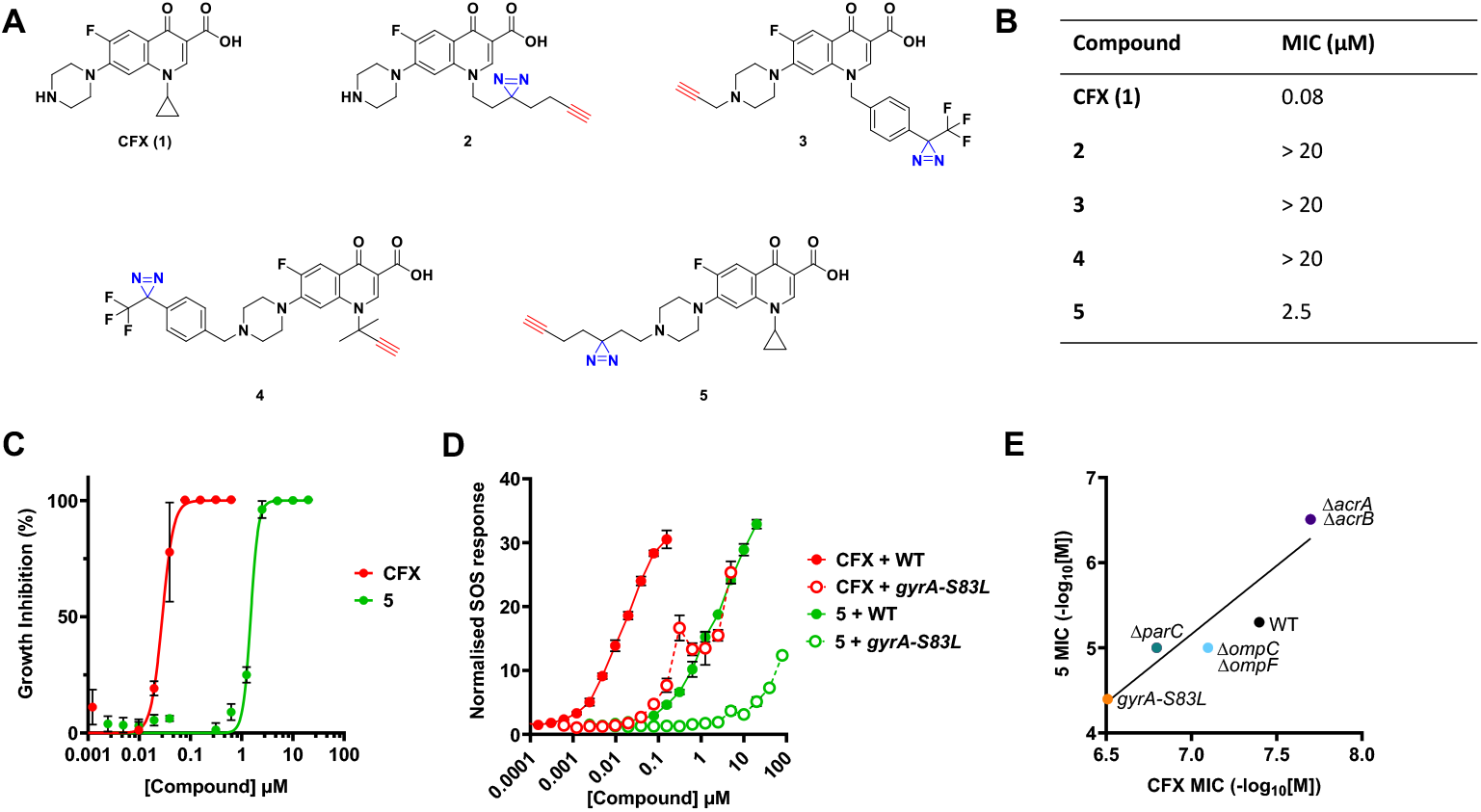
Validation of **CFX** photoaffinity probes for target profiling. A) Structures of **CFX** (**1**) and probes used in this study. B) Minimum Inhibitory Concentration (MIC) of compounds in wild-type (WT) *E. coli*. C) Dose-response of growth inhibition of WT *E. coli* by **CFX** and **5**.D) Induction of the SOS response in WT and gyrA-S83L *E. coli psulA-gfp* reporter strains, measured as GFP/OD_600_. E) Comparison of **CFX** and **5** MIC values in mutant or knockout strains known to affect **CFX** potency. Data represent mean ± standard error of the mean (SEM), n=3.

Identifying the target(s) and mechanism(s) of action (MoA) of antibiotics has traditionally exploited resistance mechanisms, by identifying either mutations or alterations in gene expression levels that correlate with resistance.^[20]^ Whilst these approaches have generated numerous invaluable insights, they may be biased towards targets with the most substantial impact on bacterial viability, overlook binding partners that do not easily tolerate mutation,^[20]^ or neglect proteins that are liganded by antibiotics but where the protein function is not inhibited. The latter class of binding partners may represent an untapped source of bacterial proteins with identified, cell-penetrant ligands that could be modified to disrupt bacterial cellular processes.

Here, we sought to complement previous resistance-based screens and uncover so far enigmatic binding partners by determining the interaction profile of **CFX** in live *E. coli* cells using affinity-based protein profiling (AfBPP) and chemical proteomics.^[21]^ AfBPP identified two previously unannotated proteins, YjdN and YbaA, as **CFX** binding partners for the first time, which were validated using genetic and biophysical approaches.

## Results and Discussion

### Design, Synthesis and Validation of CFX Probes

To comprehensively map **CFX** binding proteins in *E. coli*, a panel of four photochemical probes were designed (Fig 1A). The probes each contain a diazirine for covalent protein crosslinking upon UV irradiation, and a terminal alkyne for bioorthogonal attachment of reporter groups using copper-catalysed alkyne−azide cycloaddition (CuAAC) ‘click chemistry’.^[21]^ Alkyl and aryl diazirines and the alkyne handle were either installed on the **CFX** piperazine ring or by replacement of the cyclopropyl ring, as these positions have been shown to tolerate modification in previous SAR studies.^[22]^ Probes **2** and **3** were prepared by a Gould–Jacobs reaction of 3,4-difluoroaniline and diethyl ethoxymethylenemalonate, followed by alkylation of the fluoroquinolone core with the respective alkylhalides (Scheme S1).^[15]^ S_N_Ar with piperazine and basic ester hydrolysis afforded probe **2**, and the precursor to probe **3** which was *N*-alkylated with propargyl bromide to afford **3**. Probe **4** was synthesised from ethyl 3-(*N,N*-dimethylamino)acrylate and 1,3,4-trifluorobenzoyl chloride, with subsequent cyclisation using 2-methyl-3-butyn-2-amine to generate the fluoroquinolone subunit.^[23]^ S_N_Ar with piperazine, ester hydrolysis and *N*-alkylation with 4-[3-(trifluoromethyl)-3*H*-diazirin-3-yl]benzyl bromide afforded probe **4** (Scheme S1).^[15]^ Probe **5** was prepared via *N*-alkylation of **CFX** using a minimalist photocrosslinkable-clickable handle.^[24]^

Probes **2**-**5** were tested in *Escherichia coli* K-12 substrain MG1655 (herein referred to as *E. coli*) for growth inhibition by determining MIC values. **CFX** showed the most potent inhibition of *E. coli* growth (MIC = 0.08 µM), followed by **5** (MIC = 2.5 µM), with all other probes displaying MIC > 20 µM in wild-type (WT) *E. coli* (Fig. 1B,C). Probe **5** was therefore selected for further validation. Formation of DNA double-strand breaks from **CFX** treatment activates the SOS response and initiates expression of ‘SOS box’ genes.^[13]^ SOS activation by **CFX** and **5** was therefore tested in a cellular reporter system expressing *gfp* under the control of the SOS promoter sequence *PsulA*,^[25]^ measuring SOS induction as GFP fluorescence per OD_600_ unit (GFP/OD_600_), in *E. coli* containing either WT gyrase or resistant *gyrA-S83L*. Consistent with growth inhibition data, both compounds activated the SOS response, with **CFX** activating SOS at lower concentrations and *gyrA-S83L* mutation decreasing the activity of both compounds (Fig. 1D).

In order to confirm probe **5** shared a similar MoA with **CFX**, MIC values were determined in a panel of *E. coli* strains with mutations or knockout of genes known to affect **CFX** potency.^[26]^ The effect of mutants/knockouts on **5** MIC reflected the effect on **CFX** MIC (Fig. 1E), albeit with lower potency for **5** compared to **CFX**. *gyrA-S83L* was the least susceptible strain to both compounds, indicating the importance of the gyrase Ser83-mediated interaction. Δ*parC* increased MICs two-fold for both compounds, which was not as substantial as the *gyrA-S83L* mutation and suggested gyrase as the primary antibiotic target. Knockout of porin proteins Δ*ompC* and Δ*ompF* increased MIC for both compounds, consistent with porin-mediated cell penetration, whereas knockout of efflux pump components Δ*acrA* and Δ*acrB* reduced the MIC of both compounds (Fig. 1E). Comparison of the **CFX** and **5** MICs in all strains demonstrated a linear correlation (Pearson r = 0.93, p = 0.0021, Fig. 1E), supporting a shared MoA between the probe and parent molecule. As **5** recapitulated the activity of **CFX** against a range of mutants and in biological assays, the probe was next used in AfBPP studies to profile **CFX**-binding proteins in *E. coli*.

### Affinity-based Protein Profiling using 5

Having validated **5** as a suitable probe for **CFX** biological activity, the half-life for photoactivation of **5** by UV irradiation was determined as 2.9 min, as quantified by HPLC (Fig. S1). For target protein labelling, overnight cultures of *E. coli* were treated with either DMSO or **5** and samples UV irradiated for 12 min to induce target crosslinking. Cells were lysed, and probe-labelled proteins functionalised by CuAAC with azido-TAMRA-biotin (AzTB) for analysis (Fig. 2A). AzTB contains a TAMRA fluorophore for in-gel fluorescence visualisation and a biotin moiety for enrichment of probe-labelled proteins by avidin pulldown (Fig. S2). Following AzTB functionalisation, proteins were separated by SDS-PAGE and target proteins visualised (Fig. 2B). Protein labelling with **5** (40 µM) was dependent on UV irradiation, and labelling increased with increasing concentrations of **5**, which interestingly indicated engagement of several proteins at lower molecular weights than known **CFX** interactors (Fig. 2B). To demonstrate that labelled proteins were engaged by **CFX**, as opposed to being non-specific background labelling or only being binding partners of **5**, competition experiments were performed by pre-treating cells with **CFX** (1, 10, 100 and 1000 µM) before crosslinking with **5** (5 µM). Dose-dependent competition for protein labelling was observed for prominent protein bands at ∼35 kDa (band A), ∼20 kDa (band B) and ∼15 kDa (band C, Fig. 2B).

**Figure 2.**
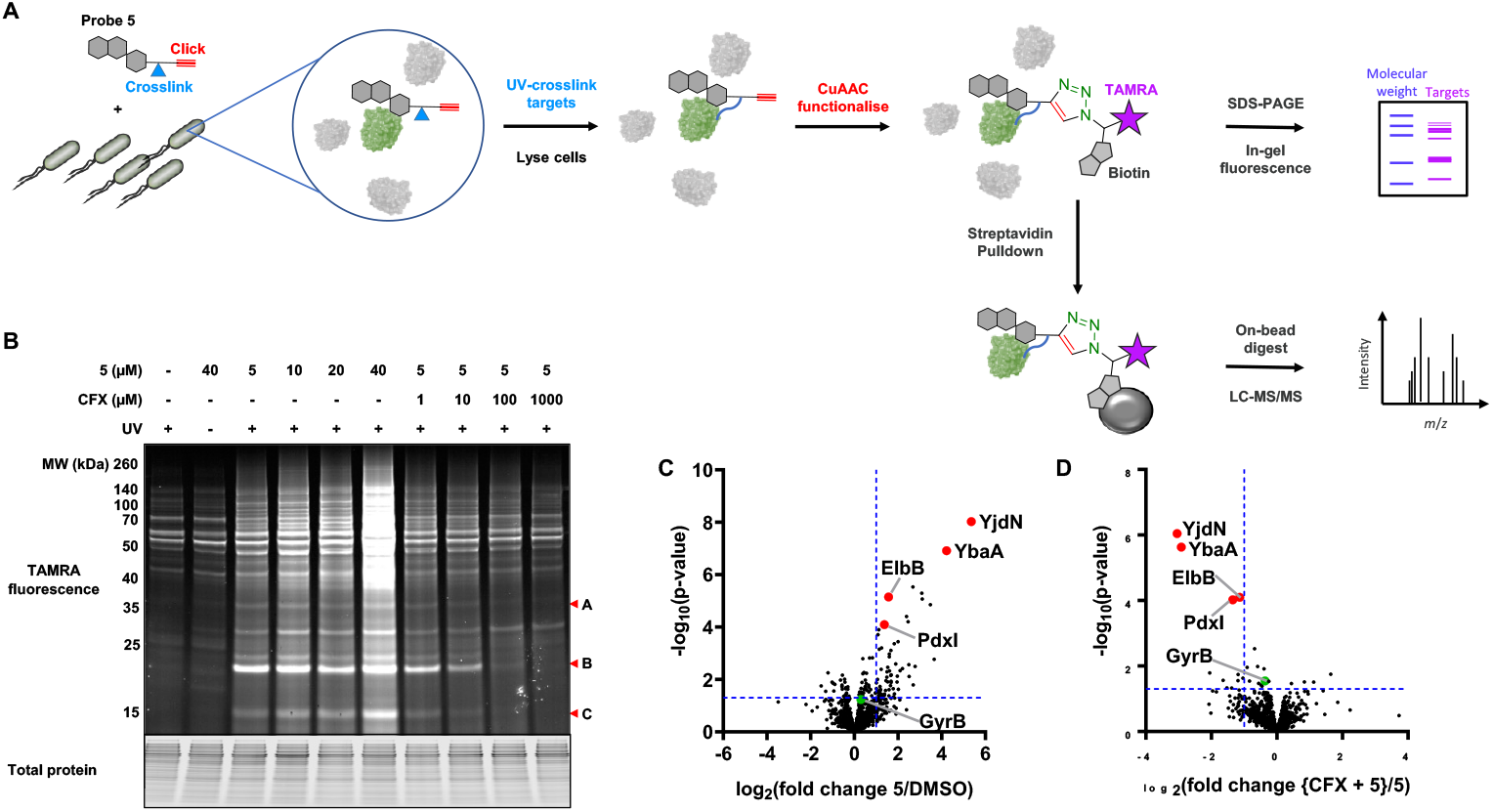
Affinity-based protein profiling (AfBPP) of **CFX** binding partners in live *E. coli*. A) AfBPP workflow. B) In-gel fluorescence of proteome labelling by **5** and **CFX** competition for protein labelling. Three prominent protein bands (A, B, C) showing **CFX** competition are annotated. Image representative of three independent experiments (n = 3). C) Proteomic analysis of protein enrichment with **5** (5 µM) compared to DMSO (n = 3). D) Proteomic analysis of protein enrichment with **5** (5 µM) in the presence of **CFX** (100 µM) compared to **5** (5 µM) alone (n = 3).

Having demonstrated that **5** successfully labelled protein binding partners of **CFX**, the identity of engaged proteins was determined by chemoproteomics. Cells were treated with DMSO, **5** (5 µM) alone, or **5** (5 µM) in competition with **CFX** (100 µM), and labelled proteins enriched using NeutrAvidin agarose (Fig. 3A). Bound proteins were washed, digested with LysC and trypsin, and the resulting peptides analysed by tandem LC-MS/MS. Comparison of DMSO- and **5**-(5 µM) treated samples identified 71 hit proteins as statistically significantly enriched by **5** (log_2_ fold-change >1 and p-value <0.05, Fig. 2C, Table S1). Comparison of competition samples treated with **CFX** (100 µM) and **5** (5 µM) against samples treated with **5** (5 µM) alone further identified 11 hit proteins competed by **CFX** with high statistical significance (log_2_ enrichment < −1 and p-value <0.05, Fig. 2D, Table S1). New target proteins were defined as hits in both conditions, with seven proteins meeting these criteria: YjdN, YbaA, ElbB, PdxI, RplW, GlpT and NupC (Table S1). YjdN, YbaA, ElbB and PdxI showed the highest statistical significance (Table 1) and were therefore selected for further investigation.

**Table 1.**
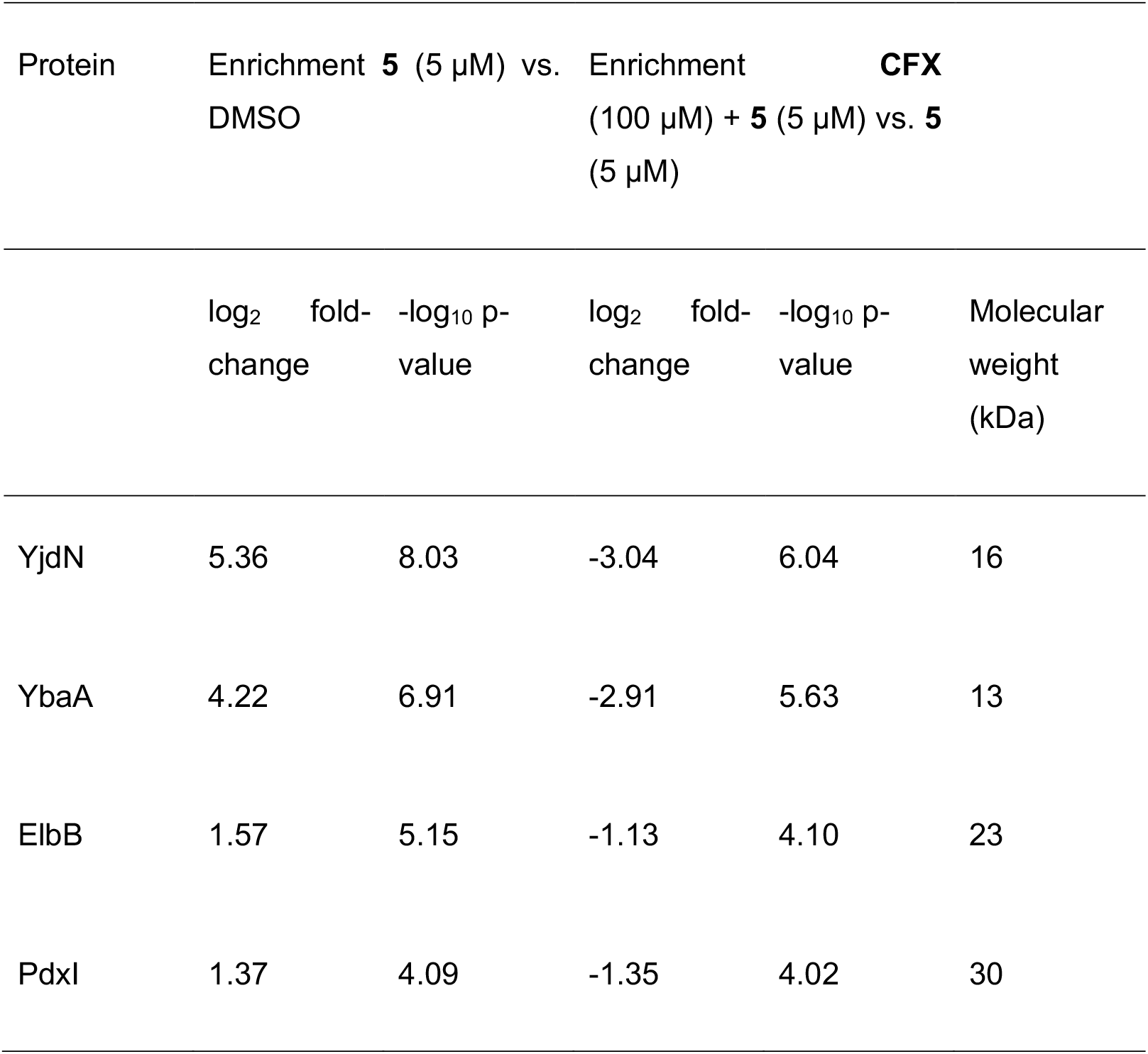
Chemoproteomic results for selected protein targets.

**Figure 3.**
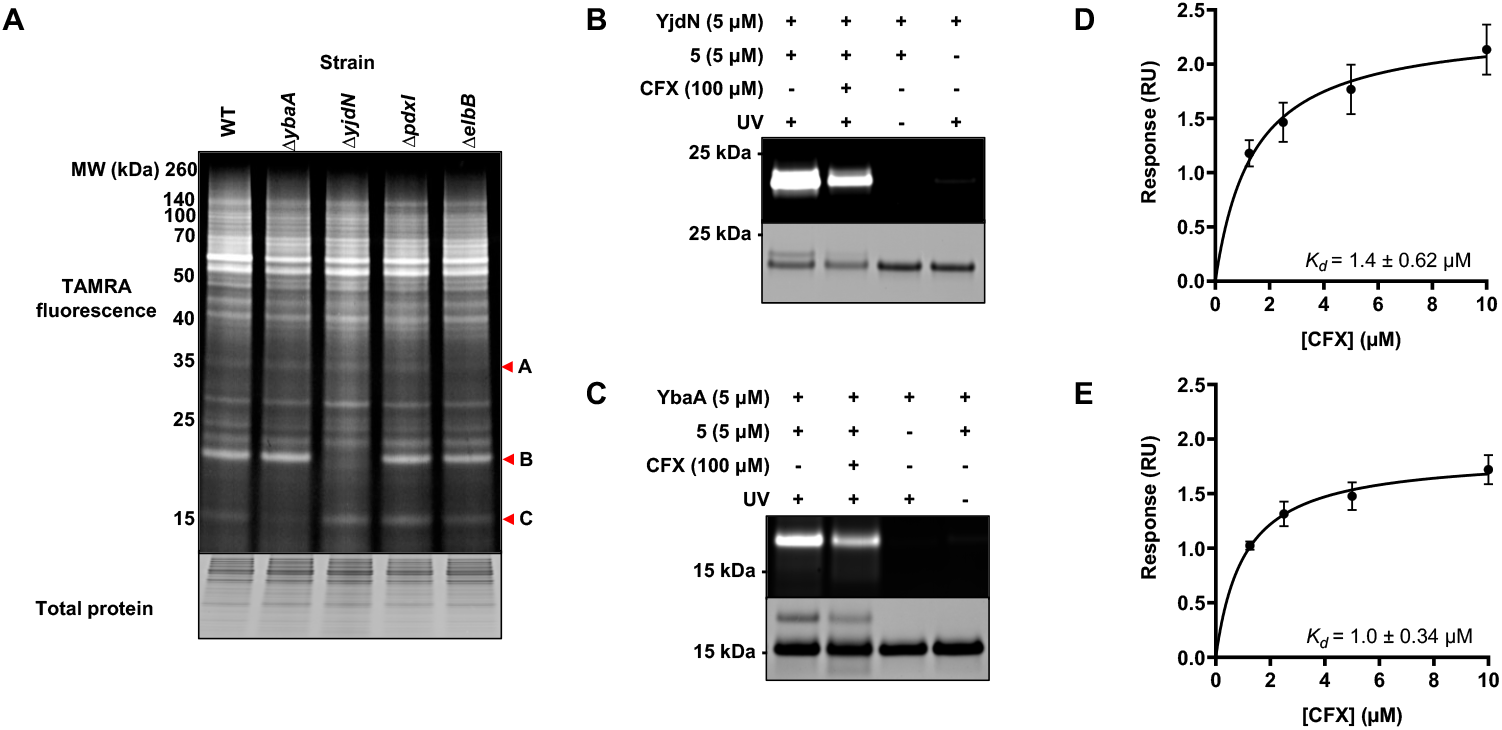
Validation of CFX binding partners. A) AfBPP in-gel fluorescence analysis of single gene knockouts of hits from chemoproteomics. B) AfBPP labelling of purified YjdN by **5** (5 µM) and competition by **CFX** (100 µM). C) AfBPP labelling of purified YbaA by **5** (5 µM) and competition by **CFX** (100 µM). D) Surface plasmon resonance of **CFX** binding to YjdN. E) Surface plasmon resonance of **CFX** binding to YbaA. Images representative of three independent experiments; data represent mean ± SEM (n = 3).

Known target GyrB was enriched by **5** (log_2_ enrichment = 0.30, p-value = 0.058) and competed by **CFX** (log_2_ enrichment = −0.36, p-value = 0.029); this lower level of enrichment of a primary **CFX** antibiotic target is consistent with the decreased toxicity and SOS activation by **5** in *E. coli* (Fig. 1). Chemoproteomics also identified known transiently-interacting proteins involved in **CFX** cell penetration and efflux (AcrB, TolC and OmpC) as enriched by **5** and competed by **CFX**, although not reaching the defined cut-off values for hit proteins (Table S2).

### Validation and Investigation of Novel CFX Binding Proteins

To validate the new **CFX**-binding proteins identified by chemoproteomics, engagement of **5** was examined by in-gel fluorescence profiling in Δ*yjdN*, Δ*ybaA*, Δ*pdxI* and Δ*elbB* knockout *E. coli* strains (Fig. 3A).^[26]^ In-gel fluorescence analysis revealed YjdN to be band B, and YbaA to be band C. No loss of labelled protein fluorescence was observed for Δ*pdxI*, whilst a decrease in the weak fluorescent labelling of the protein at ∼35 kDa was observed with Δ*elbB* knockout (Fig. 3A), suggesting ElbB may represent band A. Both PdxI and ElbB had lower enrichment with **5** in chemoproteomics (Fig. 2C), suggesting these targets may be less likely to be visualised by in-gel fluorescence.

To confirm interaction with **CFX**, YjdN and YbaA with C-terminal His_10_ tags were expressed and purified by immobilized metal affinity chromatography followed by size exclusion chromatography (Fig. S3). Purified proteins (5 µM) were photo-crosslinked with **5** (5 µM), either with or without competition from **CFX** (100 µM), functionalised by CuAAC with AzTB and visualised by in-gel fluorescence (Fig. 3B,C). Both YjdN and YbaA showed crosslinking with **5** *in vitro*, and a reduction of labelling with **CFX** competition. A ∼1 kDa increase in molecular weight was observed for both proteins following AzTB functionalisation (Fig. 3B,C), consistent with previous studies of other low molecular weight proteins.^[27]^ Direct binding of **CFX** to YjdN or YbaA was further assessed by surface plasmon resonance (SPR), which measured *K*_d_ values of 1.4 ± 0.62 µM and 1.0 ± 0.34 µM respectively (Fig. 3D,E). SPR represents a direct and probe-independent measurement of **CFX** interaction with YbaA and YjdN, and further validates use of **5** as a probe for **CFX** binders.

To further investigate the interaction of **CFX** with YjdN and YbaA, identification of the ligand binding site was attempted using **5** and proteomic sequencing. YjdN and YbaA (15 µg) were photocrosslinked to **5**, digested with LysC and trypsin, and the resulting peptides analysed by tandem LC-MS/MS.^[28]^ A total sequence coverage of 89% and 83% was achieved for YjdN and YbaA, respectively. Although a modified peptide could not be detected in YjdN, analysis of YbaA identified one peptide with an increased mass corresponding to crosslinking of **5** (IVECWASDVPDGKVTDFR), and MS/MS fragmentation identified Glu40 as the modified residue (Fig. S4). The X-ray structure of YbaA from *Shigella flexneri* has been solved (PDB: 2okq),^[29]^ which has an identical sequence to *E. coli* YbaA. The structure indicates YbaA adopts an α-β barrel conformation with a central pocket at the α-β interface; Glu40 is located at the bottom of the pocket, therefore suggesting a potential ligand-binding site (Fig. 4A,B). To confirm the importance of Glu40 for **CFX** interaction with YbaA, Glu40Lys and Glu40Ala mutants were expressed and purified (Fig. S5). *In vitro* crosslinking with **5** demonstrated a loss of YbaA interaction with either mutant (Fig. 4C). Binding of **5** in proximity to the negatively-charged Glu40 suggested the potential for an ionic or hydrogen-bonding interaction with the positively charged piperidine nitrogen of **CFX** (Fig S6); however, the diazo intermediate formed by UV activation of alkyl diazirines is also known to preferentially react with acidic residues such as glutamate and aspartate.^[30]^ A **CFX** analogue replacing the piperazine ring with a dimethylamino group (**6**, Fig. 4D) was therefore synthesised (Scheme S2) to test for a putative interaction. AfBPP with **5** demonstrated decreased competition for YbaA binding with **6** (Fig. 4E), supporting the existence of an interaction between the piperidine nitrogen of **CFX** and Glu40 of YbaA in the binding pocket.

**Figure 4.**
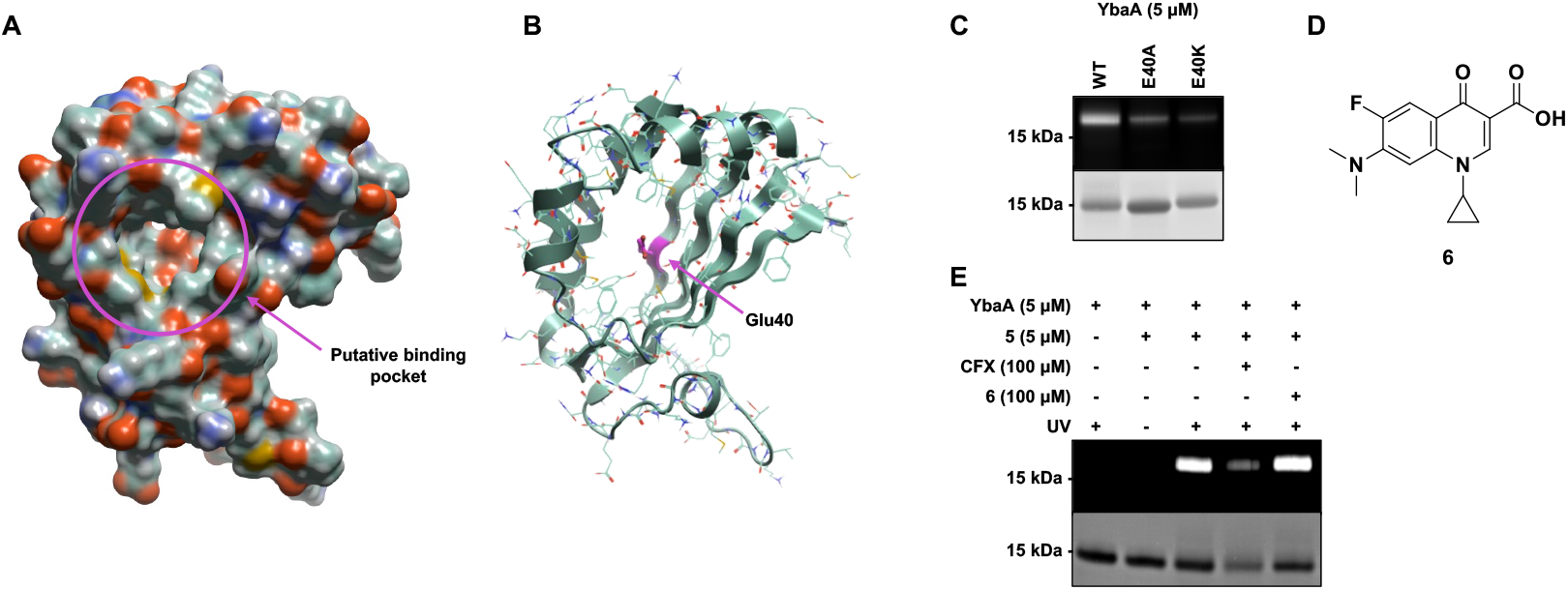
**CFX**-binding site identification in YbaA. A) Surface view of *S. flexneri* YbaA (PDB: 2okq)^[29]^, which is sequence identical to *E. coli* YbaA, with putative binding pocket indicated (purple). B) YbaA cartoon representation with Glu40 residue modified by **5** displayed in purple. C) Mutation of YbaA-His_10_ Glu 40 to Ala or Lys results in decreased labelling by **5** (5 µM). D) **CFX** analogue **6** with piperazine ring exchanged to dimethylamine. E) Removal of ionisable nitrogen results in decreased competition from analogue **6** (100 µM) for YbaA-His_10_ labelling by probe **5** (5 µM). Images representative of three independent experiments (n = 3).

The functions of YjdN and YbaA are currently unknown. *yjdN* (also named *phnB*) is located in the *phn* locus that encodes proteins necessary for *E. coli* growth with alkylphosphonates as the sole phosphorous source;^[31]^ however, subsequent studies indicate YjdN has no role in phosphonate metabolism.^[32]^ YbaA is unannotated except for a domain of unknown function (DUF1428) of which many family members are annotated as a 4.5S RNA signal recognition particle, although this remains to be verified (Pfam entry PF07237).^[33]^ To investigate possible effects from **CFX** binding to YjdN and YbaA in cells, the **CFX** MIC and SOS response activation were measured in knockout *E. coli* strains. However, no effect on **CFX** potency or SOS activation was observed from target knockout (Fig. S7). The *K*_d_ values for binding to YjdN or YbaA in SPR are ∼1 μM and therefore above the MIC of **CFX** in WT *E. coli* (∼80 nM), which may explain the lack of effect from knockout in these assays. The effect of **CFX** interaction with YbaA and YjdN in other biological pathways not essential for cell growth *in vitro* remains to be determined.

## Conclusion

Fluoroquinolones such as **CFX** are critical antibiotic agents for a range of infections, yet various aspects of the MoA and susceptibility to resistance development of **CFX** are not fully understood despite over 40 years of clinical use.^[5, 19]^. We present here a study of **CFX**-binding proteins in live *E. coli* using target-agnostic AfBPP. Probe **5** was selected for target profiling studies as it recapitulated **CFX** biological activity in a range of mutant strains and assays, albeit with lower potency. During preparation of this manuscript, a chemoproteomic study of **CFX** off-targets in human cells independently developed probe **5** and demonstrated ∼16-fold higher IC_50_ for ParCE inhibition *in vitro*;^[34]^ the decreased toxicity and SOS activation of **5** may therefore result from weaker inhibition of the primary topoisomerase antibiotic targets. AfBPP revealed previously unreported **CFX**-binding partners, YbaA and YjdN, that were validated using genetic and biophysical approaches. The use of photochemical probes and chemical proteomics further provided insights into key interactions in the **CFX**-binding site of YbaA. The functions of YbaA and YjdN are currently unknown, as is the effect of **CFX** binding to these proteins. The ∼1 μM *K*_d_ for interaction of **CFX** with these proteins suggests that engagement may be more substantial in **CFX**-resistant strains containing mutant DNA gyrase and topoisomerase IV. In these strains, **CFX** binding to YjdN/YbaA may produce phenotypes that are beneficial to bacterial survival or alternatively promote bacterial killing/clearance, and is the subject of on-going investigations. Collectively, the full target profile of **CFX** in *E. coli* and associated tools and methods represent a means to generate deeper understanding of existing antibiotic MoAs and resistance to support future studies.

## Supporting Information

The authors have cited additional references within the Supporting Information.^[35-37]^

## Acknowledgements

This research was supported by the Wellcome Trust (Grant Number: 317713/Z/24/Z and 218514/Z/19/Z), the Ineos Oxford Institute for Antimicrobial Research, and a Springboard Award from the Academy of Medical Sciences (AMS), the Wellcome Trust, the Government Department of Business, Energy and Industrial Strategy (BEIS), the British Heart Foundation and Diabetes UK [SBF007\100164]. The authors thank Dr Vaishnavi Ravikumar and Dr. Marjorie Fournier (Advanced Proteomics Facility, Department of Biochemistry, University of Oxford) for assistance with proteomic experiments and analysis.

## Data Availability Statement

The mass spectrometry proteomics data have been deposited to the ProteomeXchange Consortium via the PRIDE partner repository with the dataset identifier PXD068697.

## References

[1] Antimicrobial Resistance Collaborators, Lancet 2022, 399, 629–655.

[2] K. Allel, L. Day, A. Hamilton, L. Lin, L. Furuya-Kanamori, C. E. Moore, T. Van Boeckel, R. Laxminarayan, L. Yakob, Lancet Planet Health 2023, 7, e291–e303.

[3] D. Chinemerem Nwobodo, M. C. Ugwu, C. Oliseloke Anie, M. T. S. Al-Ouqaili, J. Chinedu Ikem, U. Victor Chigozie, M. Saki, J Clin Lab Anal 2022, 36, e24655.

[4] WHO, 2023, WHO/MHP/HPS/EML/2023.2002.

[5] A. Shariati, M. Arshadi, M. A. Khosrojerdi, M. Abedinzadeh, M. Ganjalishahi, A. Maleki, M. Heidary, S. Khoshnood, Front Public Health 2022, 10, 1025633.

[6] N. G. Bush, K. Evans-Roberts, A. Maxwell, EcoSal Plus 2015, 6.

[7] A. B. Khodursky, E. L. Zechiedrich, N. R. Cozzarelli, Proc. Natl. Acad. Sci. U.S.A. 1995, 92, 11801–11805.

[8] S. Bagel, V. Hullen, B. Wiedemann, P. Heisig, Antimicrob Agents Chemother 1999, 43, 868–875.

[9] G. A. Jacoby, Clin. Infect. Dis. 2005, 41 Suppl 2, S120–126.

[10] G. Dong, J. Li, L. Chen, W. Bi, X. Zhang, H. Liu, X. Zhi, T. Zhou, J. Cao, Braz J Infect Dis 2019, 23, 15–21.

[11] Z. Rafaque, N. Abid, N. Liaqat, P. Afridi, S. Siddique, S. Masood, S. Kanwal, J. Dasti, Infect Drug Resist 2020, 13, 2801–2810.

[12] S. Whelan, M. C. O’Grady, G. D. Corcoran, K. Finn, B. Lucey, Med Sci (Basel) 2022, 11.

[13] T. Lanyon-Hogg, Future Med Chem 2021, 13, 143–155.

[14] A. Winnifrith, S. R. Brown, P. Jedryszek, C. Grant, P. E. Kay, A. M. Thomas, J. D. Bradbury, T. Lanyon-Hogg, RSC Chem Biol 2025, 6, 772–779.

[15] J. D. Bradbury, T. Hodgkinson, A. M. Thomas, O. Tanwar, G. La Monica, V. V. Rogga, L. J. Mackay, E. K. Taylor, K. Gilbert, Y. Zhu, A. Y. Sefton, A. M. Edwards, C. J. Gray-Hammerton, G. R. Smith, P. M. Roberts, T. R. Walsh, T. Lanyon-Hogg, Chem Sci 2024, 15, 9620–9629.

[16] J. L. Gray, E. V. K. Ledger, T. Suwatthee, T. J. Burden, K. Arvaniti, A. Sefton, L. E. Papagora, T. B. Clarke, J. Riley, E. G. Pinto, F. Cunningham, I. H. Gilbert, D. Gray, D. N. Wang, K. D. Read, T. Lanyon-Hogg, N. J. Traaseth, A. M. Edwards, E. W. Tate, bioRxiv 2025.

[17] C. S. Q. Lim, K. P. Ha, R. S. Clarke, L. A. Gavin, D. T. Cook, J. A. Hutton, C. L. Sutherell, A. M. Edwards, L. E. Evans, E. W. Tate, T. Lanyon-Hogg, Bioorg. Med. Chem. 2019, 27, 114962.

[18] S. K. Morgan-Linnell, L. Becnel Boyd, D. Steffen, L. Zechiedrich, Antimicrob Agents Chemother 2009, 53, 235–241.

[19] J. Ruiz, Life (Basel) 2024, 14.

[20] M. A. Hudson, S. W. Lockless, mBio 2022, 13, e0224021.

[21] H. Fang, B. Peng, S. Y. Ong, Q. Wu, L. Li, S. Q. Yao, Chem Sci 2021, 12, 8288–8310.

[22] L. R. Peterson, Clin. Infect. Dis. 2001, 33 Suppl 3, S180–186.

[23] R. A. Bunce, E. J. Lee, M. T. Grant, J. Heterocycl. Chem. 2011, 48, 620–625.

[24] Z. Li, P. Hao, L. Li, C. Y. Tan, X. Cheng, G. Y. Chen, S. K. Sze, H. M. Shen, S. Q. Yao, Angew. Chem. Int. Ed. Engl. 2013, 52, 8551–8556.

[25] Y. Y. Cheng, Z. Zhou, J. M. Papadopoulos, J. D. Zuke, T. G. Falbel, K. Anantharaman, B. M. Burton, O. S. Venturelli, Mol. Syst. Biol. 2023, 19, e11406.

[26] T. Baba, T. Ara, M. Hasegawa, Y. Takai, Y. Okumura, M. Baba, K. A. Datsenko, M. Tomita, B. L. Wanner, H. Mori, Mol. Syst. Biol. 2006, 2, 2006 0008.

[27] T. Lanyon-Hogg, M. Ritzefeld, L. Sefer, J. K. Bickel, A. F. Rudolf, N. Panyain, G. Bineva-Todd, C. A. Ocasio, N. O’Reilly, C. Siebold, A. I. Magee, E. W. Tate, Chem Sci 2019, 10, 8995–9000.

[28] T. Lanyon-Hogg, M. Ritzefeld, L. Zhang, S. A. Andrei, B. Pogranyi, M. Mondal, L. Sefer, C. D. Johnston, C. E. Coupland, J. L. Greenfield, J. Newington, M. J. Fuchter, A. I. Magee, C. Siebold, E. W. Tate, Angew. Chem. Int. Ed. Engl. 2021, 60, 13542–13547.

[29] G. Minasov, I.I. Vorontsov, L. Shuvalova, J.S. Brunzelle, O. Kiryukhina, F.R. Collart, A. Joachimiak, W.F. Anderson, 2007, Crystal structure of unknown conserved ybaA protein from Shigella flexneri, PDB: 2okq, 10.2210/pdb2OKQ/pdb

[30] A. V. West, G. Muncipinto, H. Y. Wu, A. C. Huang, M. T. Labenski, L. H. Jones, C. M. Woo, J. Am. Chem. Soc. 2021, 143, 6691–6700.

[31] C. M. Chen, Q. Z. Ye, Z. M. Zhu, B. L. Wanner, C. T. Walsh, J. Biol. Chem. 1990, 265, 4461–4471.

[32] W. W. Metcalf, B. L. Wanner, Gene 1993, 129, 27–32.

[33] J. Mistry, S. Chuguransky, L. Williams, M. Qureshi, G. A. Salazar, E. L. L. Sonnhammer, S. C. E. Tosatto, L. Paladin, S. Raj, L. J. Richardson, R. D. Finn, A. Bateman, Nucleic Acids Res. 2021, 49, D412–D419.

[34] T. Reinhardt, Y. El Harraoui, A. Rothemann, A. T. Jauch, S. Muller-Deubert, M. F. Kollen, T. Risch, L. J. Jacobs, R. Muller, F. R. Traube, D. Docheva, S. Zahler, J. Riemer, N. C. Bach, S. A. Sieber, Angew. Chem. Int. Ed. Engl. 2025, 64, e202421424.

[35] A. T. Kong, F. V. Leprevost, D. M. Avtonomov, D. Mellacheruvu, A. I. Nesvizhskii, Nat. Methods 2017, 14, 513–520.

[36] K. M. Orritt, L. Feng, J. F. Newell, J. N. Sutton, S. Grossman, T. Germe, L. R. Abbott, H. L. Jackson, B. K. L. Bury, A. Maxwell, M. J. McPhillie, C. W. G. Fishwick, RSC Med Chem 2022, 13, 831–839.

[37] M. E. Ritchie, B. Phipson, D. Wu, Y. Hu, C. W. Law, W. Shi, G. K. Smyth, Nucleic Acids Res. 2015, 43, e47.

