## Supplementary Material for "Live-Cell Chemoproteomic Profiling Identifies the Uncharacterised Protein YbaA as a Direct Target of Ciprofloxacin in *Escherichia coli*"

### Supplementary Schemes

**A**

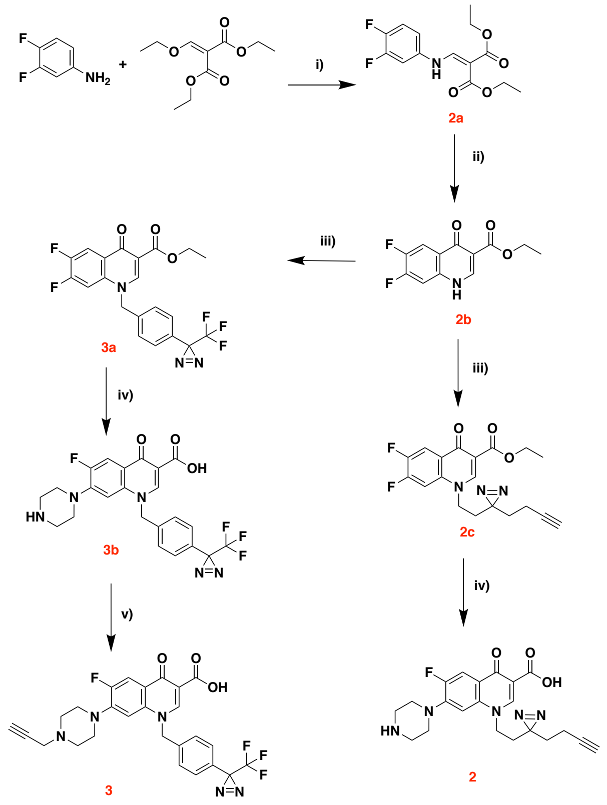

**B**

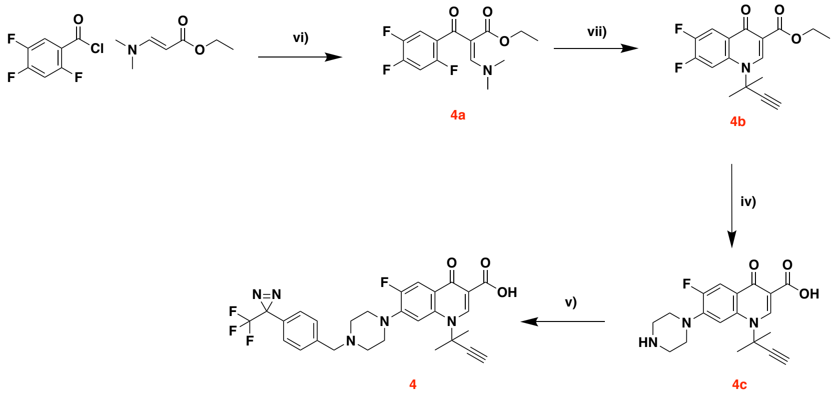

**C**

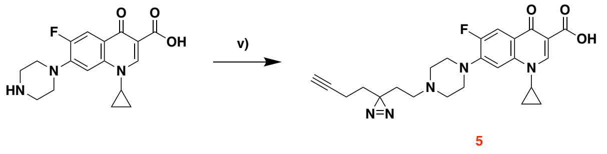

**Supplementary Scheme 1. Synthesis of CFX photochemical probes. A**) Probes **2** and **3**. **B**) Probe **4**. **C**) Probe **5**. General Reagents and Conditions: **i)** 120 °C, 18 h. **ii)** 250 °C, 1 h. **iii)** R-X, $K_2CO_3$, DMF, RT, 18 h. **iv)** a) Piperazine, MeCN, 50 °C, 18 h. b)

aq. 1 M NaOH, 50 °C, 2 h. **v)** R-X, $K_2CO_3$, MeCN, RT, 18 h. **vi)** a) $Et_2O$:EtOH (1:1), RT, 4 h. b) $K_2CO_3$, DMF, 100 °C, 4 h. **vii)** a) 2-methyl-3-butyn-2-amine, $Et_2O$, EtOH, RT, 4 h. b) $K_2CO_3$, DMF, 100 °C, 4 h.

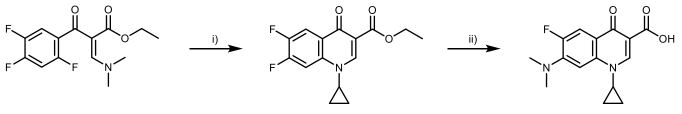

**Supplementary Scheme 2. Synthesis of CFX analogue 6.** Reagents and Conditions: **i)** a) cyclopropylamine, $Et_2O$, EtOH, RT, 3 h. (b) $K_2CO_3$, DMF, 100 °C, 18 h. **ii)** a) Dimethylamine, MeCN, 50 °C, 16 h. b) aq. 1 M NaOH, 50 °C, 2 h.

### Supplementary Figures

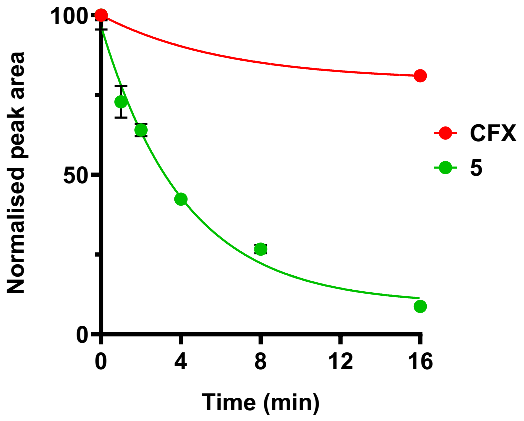

**Figure S1 – Photoactivation of 5 determined by HPLC.** Data represent mean ± standard error of the mean (SEM, n = 2).

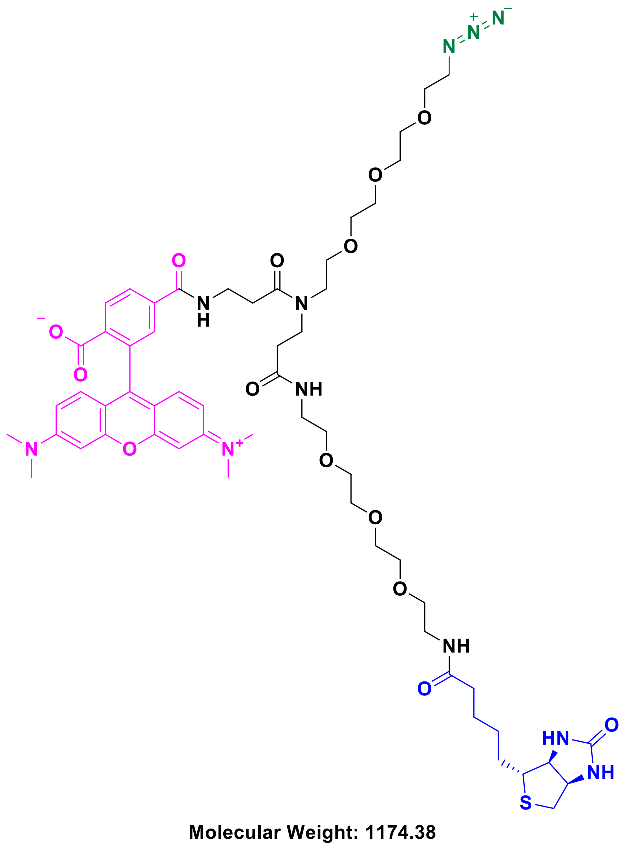

**Figure S2. Structure of azide-TAMRA-biotin (AzTB) capture reagent.** Key functional groups in AzTB: azide (green) for copper(I)-catalysed azide-alkyne cycloaddition (CuAAC); TAMRA (pink) for in-gel fluorescence visualisation; biotin (blue) for enrichment of labelled proteins.

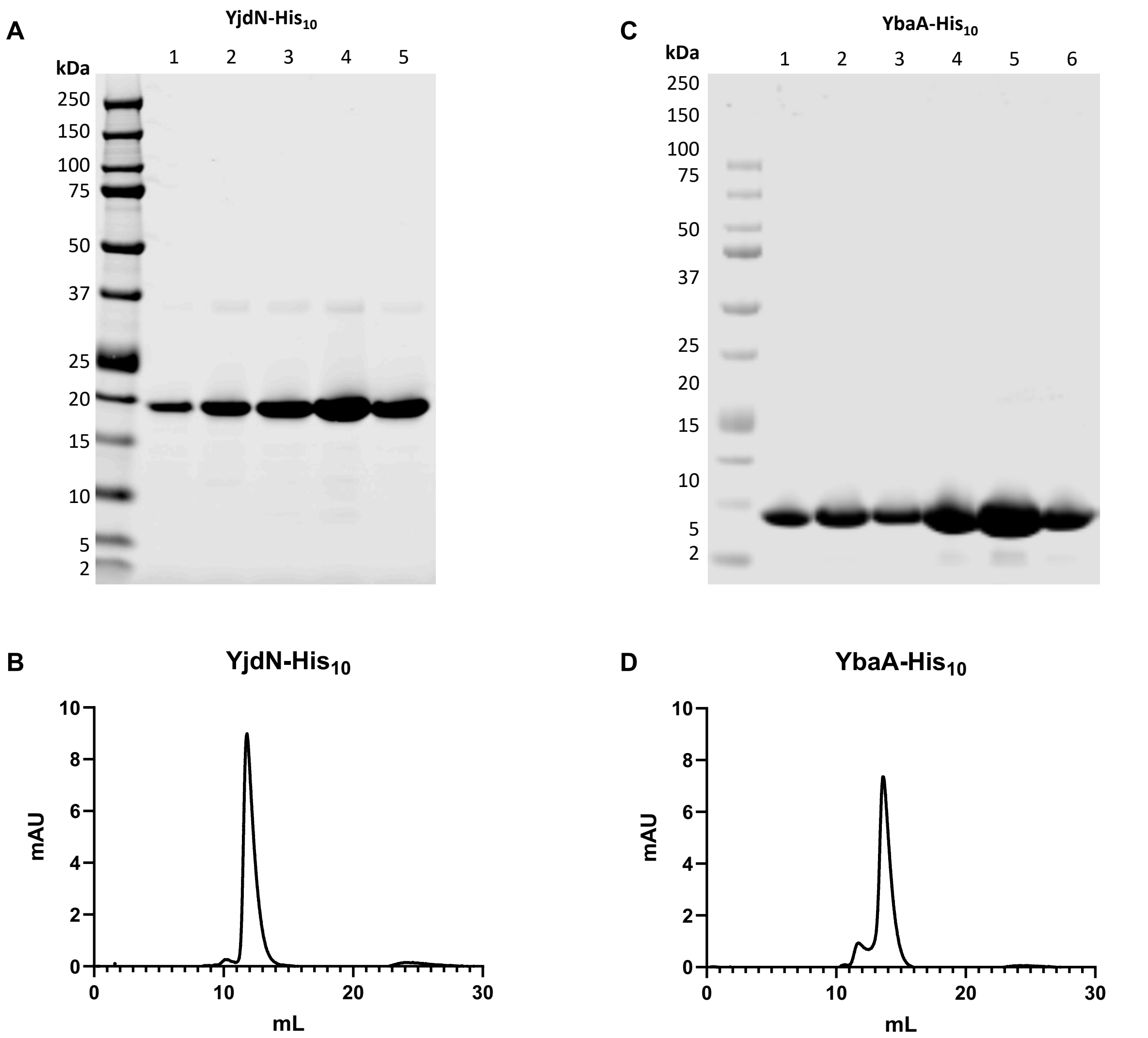

**Figure S3. Purification of recombinant proteins. A**) SDS-PAGE analysis of YjdN-$His_{10}$ size-exclusion chromatography (SEC) purification. **B**) Analytical SEC trace of pooled YjdN-$His_{10}$ fractions 4 and 5. **C**) SDS-PAGE analysis of YbaA-$His_{10}$ SEC purification. **D**) Analytical SEC trace of pooled YbaA-$His_{10}$ fractions 5 and 6.

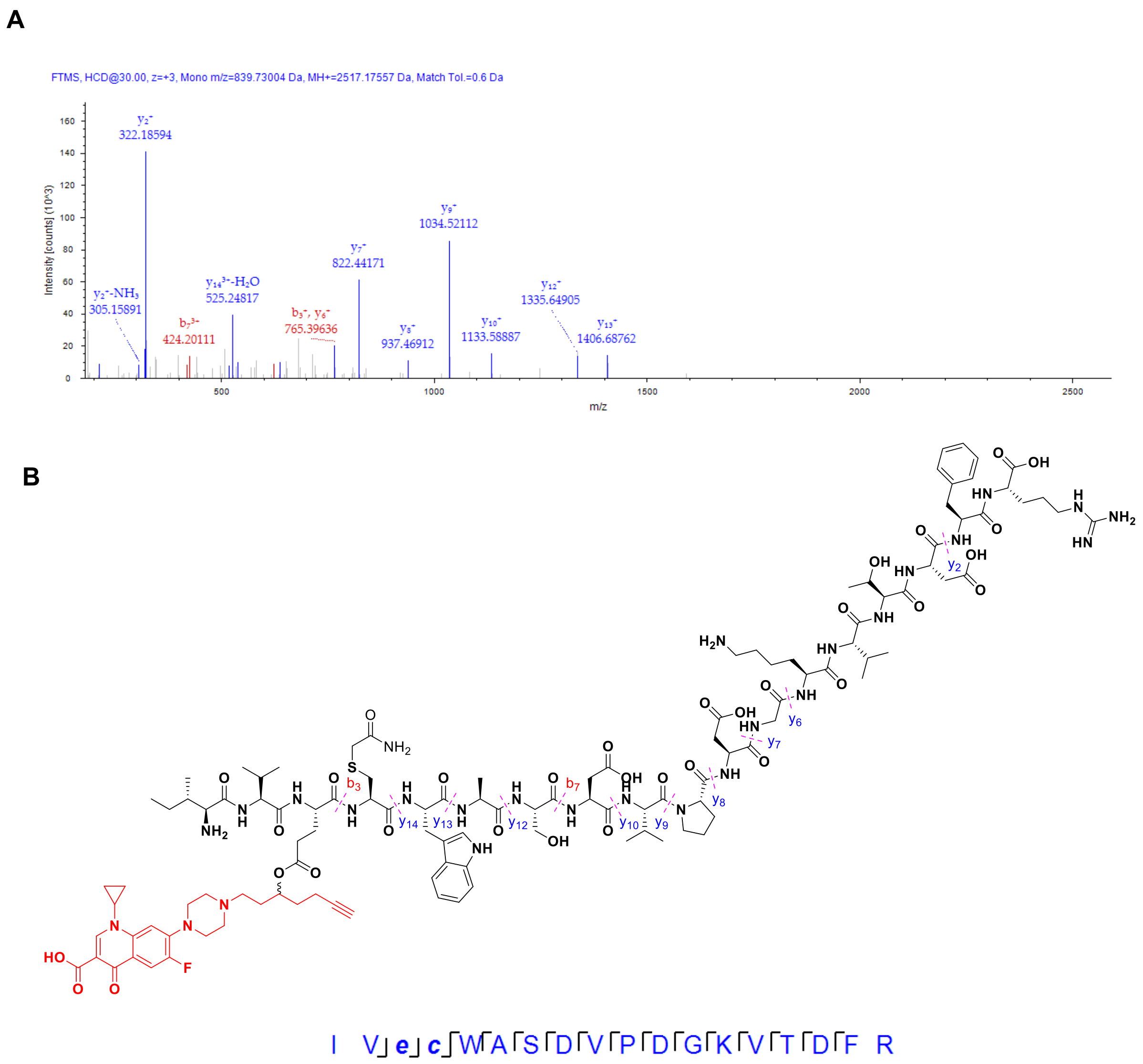

**Figure S4. LC-MS/MS analysis of YbaA-His$_{10}$ peptide labelled by probe 5. A**) LC-MS/MS spectra for YbaA peptide IVECWASDVPDGKVTDFR (PEP = $2.74 \times 10^{-13}$). **B**) Structure of peptide modified with probe **5** and fragmentation sites.

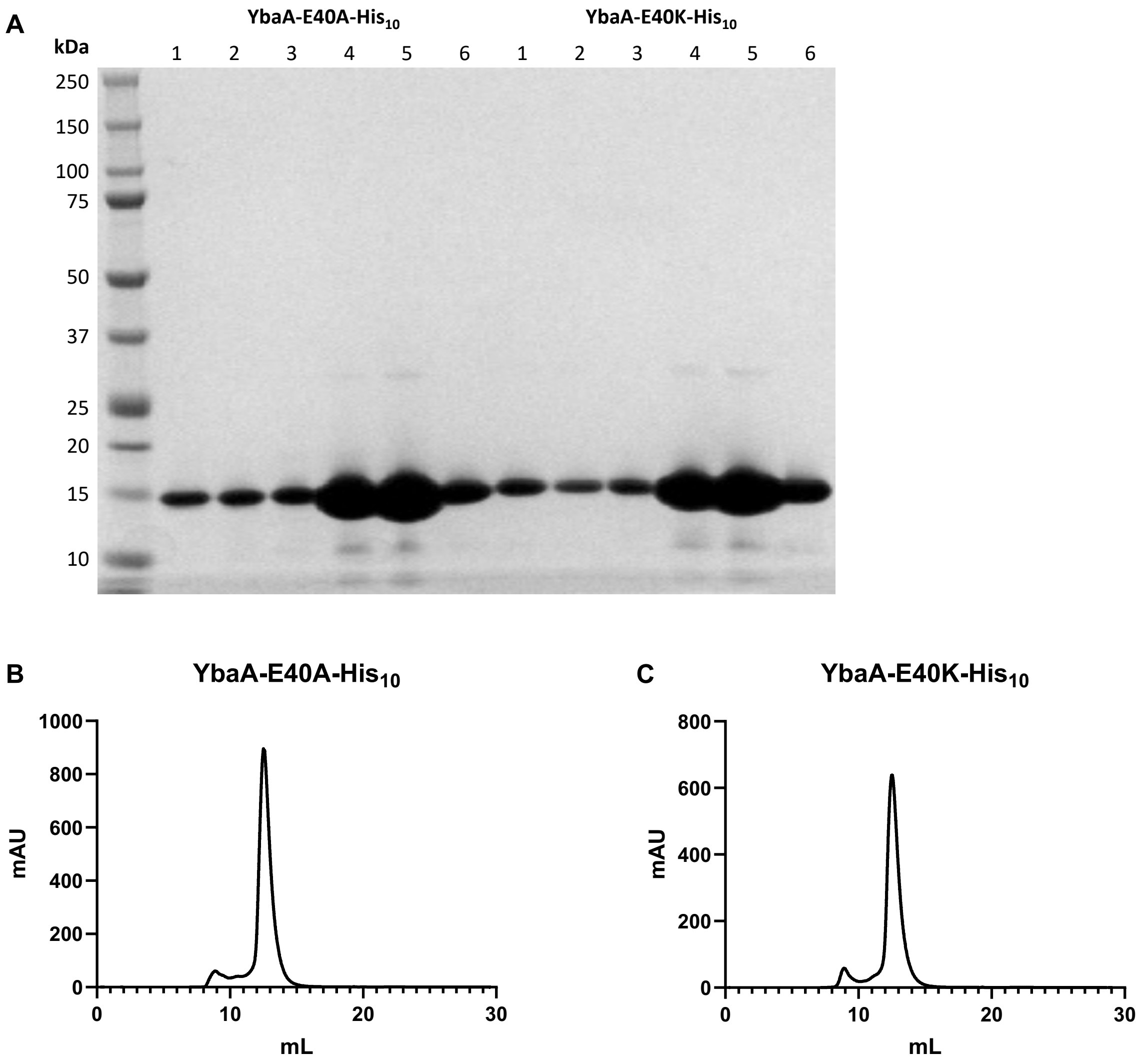

**Figure S5. Purification of YbaA mutants. A**) SDS-PAGE analysis of YbaA-E40A-$His_{10}$ and YbaA-E40K-$His_{10}$ SEC purification. **B**) Analytical SEC trace of pooled YbaA-E40A-$His_{10}$ fractions 4, 5 and 6. **C**) Analytical SEC trace of pooled YbaA-E40K-$His_{10}$ fractions 4, 5 and 6.

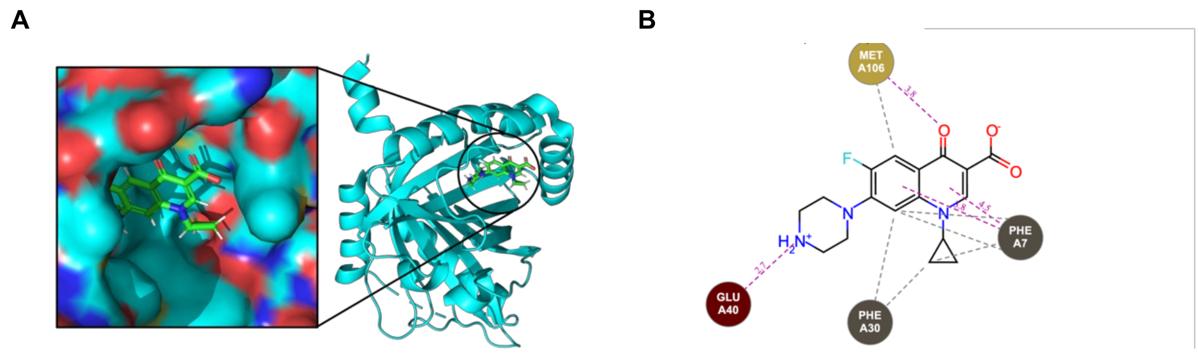

**Figure S6. *In silico* prediction of YbaA-CFX interaction. A**) Docking of **CFX** into *S. flexneri* YbaA structure (PDB: 2okq). *S. flexneri* YbaA is sequence identical to *E. coli* YbaA. **B**) Predicted interaction map between **CFX** and YbaA, suggesting the presence of an ionic interaction between the positively-charged piperazine nitrogen and negatively-charged Glu40 residue modified by probe **5**.

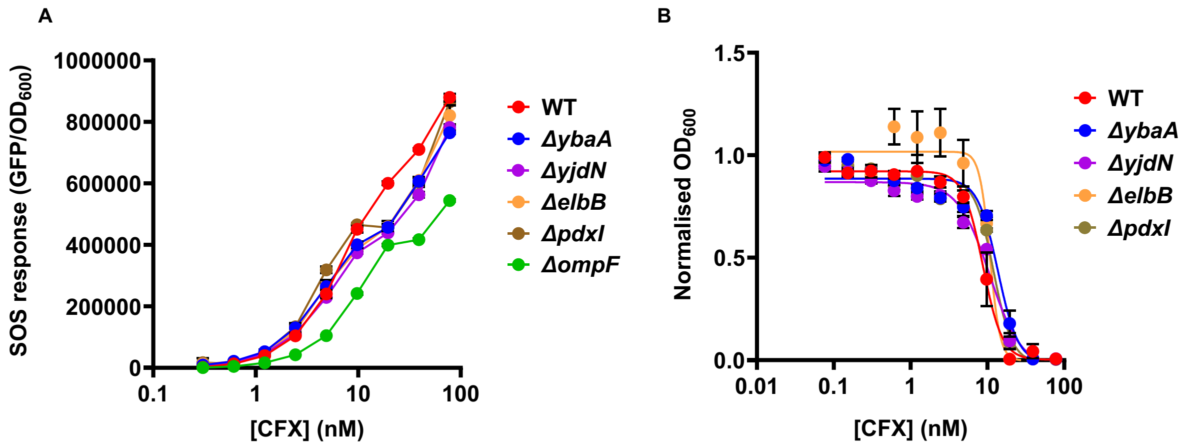

**Figure S7. Effect of newly-identified targets on SOS induction and CFX potency.** **A**) Induction of the SOS response *E. coli* target knockout *strains* transformed with the *psulA-gfp* reporter plasmid, measured as GFP/$OD_{600}$. The Δ*ompF* strain was included as a control to demonstrate the effect of knockout of a target shown to affect **CFX** potency (Fig. 1E), indicating no substantial effect from knockout of newly-identified targets. **B**) Comparison of **CFX** potency in knockout strains, demonstrating no substantial effect from knockout of newly-identified targets. Data represent mean ± SEM (n = 3).

**Supplementary Tables**

**Table S1. List of all protein hits from either probe enrichment or CFX competition chemoproteomics (n = 3).** Hit proteins were defined as those with $log_2$(fold-change) >1 and p-value <0.05 for enrichment with **5** (5 µM, Fig 2C), or $log_2$(fold-change) <-1 and p-value <0.05 for competition with **CFX** (100 µM, Fig 2D). Target proteins were defined as hits in both conditions (green).

| | | **5** vs DMSO | | | (**5**+ CFX) vs **5** | | |
|---|---|---|---|---|---|---|---|
| Protein.ID | Gene | $log_2$(fold-change) | p-value | Hit? | $log_2$(fold-change) | p-value | Hit? |
| P16681 | yjdN | 5.35573 | 9.42E-09 | Y | -3.04539 | 9.12E-07 | Y |
| P0AAQ6 | ybaA | 4.224106 | 1.22E-07 | Y | -2.91622 | 2.35E-06 | Y |
| P0ABU5 | elbB | 1.567655 | 7.14E-06 | Y | -1.13683 | 7.99E-05 | Y |
| P25906 | pdxI | 1.379987 | 8.08E-05 | Y | -1.34877 | 9.54E-05 | Y |
| P76569 | yfgD | 0.553661 | 0.440677 | | -2.05132 | 0.016333 | Y |
| P0ADZ0 | rplW | 1.069263 | 0.028332 | Y | -1.18421 | 0.018106 | Y |
| P77774 | bamB | 0.439623 | 0.27367 | | -1.03811 | 0.023499 | Y |
| P08194 | glpT | 1.954053 | 0.012469 | Y | -1.64227 | 0.027505 | Y |
| P0AFF2 | nupC | 1.899247 | 0.002571 | Y | -1.18748 | 0.028047 | Y |
| P37636 | mdtE | -0.32811 | 0.629232 | | -1.52698 | 0.046795 | Y |
| P61175 | rplV | 0.129294 | 0.831444 | | -1.37839 | 0.047037 | Y |

| | | | | | | |
|---|---|---|---|---|---|---|
| P37648 | yhjJ | 1.272551 | 0.037845 | Y | -1.15339 | 0.05459 |
| P33012 | sbmC | 2.694484 | 0.016123 | Y | -1.96208 | 0.057941 |
| P0AGA2 | secY | 3.110178 | 8.49E-06 | Y | -0.68304 | 0.065822 |
| P0AF98 | lptF | 1.384391 | 0.039415 | Y | -1.1508 | 0.075644 |
| P02358 | rpsF | 2.013669 | 0.0237 | Y | -1.44024 | 0.082241 |
| P23871 | hemH | 2.206035 | 0.003833 | Y | -1.09866 | 0.082336 |
| P60560 | guaC | 2.780668 | 0.003184 | Y | -1.27677 | 0.095442 |
| P0A8U6 | metJ | 2.096372 | 0.004636 | Y | -1.01094 | 0.100241 |
| P0A9M2 | hpt | 1.206018 | 0.024502 | Y | -0.75156 | 0.124546 |
| P56580 | srlE | 1.385693 | 0.036136 | Y | -0.94798 | 0.124739 |
| P23843 | oppA | 1.611628 | 0.004751 | Y | -0.68624 | 0.139577 |
| P31224 | acrB | 1.009884 | 0.013468 | Y | -0.503 | 0.156856 |
| P27848 | yigL | 1.452665 | 0.011188 | Y | -0.65349 | 0.181082 |
| P0AG14 | sohB | 1.150916 | 0.04391 | Y | -0.69233 | 0.189963 |
| P77667 | sufA | 1.495052 | 0.005132 | Y | -0.5519 | 0.200706 |
| P75960 | cobB | 1.971736 | 0.007493 | Y | -0.77871 | 0.201797 |

| | | | | | | |
|---|---|---|---|---|---|---|
| P25738 | msyB | 1.245797 | 0.047992 | Y | -0.72167 | 0.215094 |
| P0AEM4 | flgM | 1.590038 | 0.024553 | Y | -0.71423 | 0.252123 |
| P31550 | thiB | 1.6854 | 0.011779 | Y | -0.63483 | 0.259707 |
| P37340 | mdtK | 2.507153 | 0.007651 | Y | -0.83008 | 0.279329 |
| P29744 | flgL | 1.683636 | 0.012965 | Y | -0.60914 | 0.286341 |
| P0A9H7 | cfa | 1.194656 | 0.02553 | Y | -0.49589 | 0.291226 |
| P0ACD4 | iscU | 2.390887 | 3.95E-05 | Y | 0.32188 | 0.319195 |
| P0AEX9 | malE | 2.499007 | 0.002729 | Y | -0.60527 | 0.33169 |
| P60716 | lipA | 1.109592 | 0.041009 | Y | -0.45769 | 0.34715 |
| P77804 | ydgA | 2.686206 | 2.94E-06 | Y | -0.23244 | 0.366019 |
| P76002 | pliG | 1.655175 | 0.017241 | Y | -0.51204 | 0.384824 |
| P0A6A8 | acpP | 1.124511 | 0.000126 | Y | -0.1445 | 0.412823 |
| P0A7X6 | rimM | 1.143503 | 0.047353 | Y | -0.39915 | 0.439654 |
| P0AFK9 | potD | 1.688266 | 0.006667 | Y | -0.3668 | 0.457346 |
| P25553 | aldA | 1.143351 | 0.000707 | Y | -0.15459 | 0.49952 |
| P20605 | fic | 1.604997 | 0.001069 | Y | 0.224448 | 0.512044 |

| | | | | | | |
|---|---|---|---|---|---|---|
| P39176 | erfK | 1.603987 | 0.001371 | Y | -0.23154 | 0.515194 |
| A0A377CXR8 | pepN_1 | 3.087272 | 5.11E-06 | Y | -0.19882 | 0.525314 |
| P63020 | nfuA | 1.531033 | 0.012336 | Y | -0.30943 | 0.537597 |
| P32162 | yiiS | 1.83444 | 0.035519 | Y | -0.42986 | 0.572617 |
| P07001 | pntA | 1.081333 | 0.000196 | Y | -0.09322 | 0.600885 |
| P76142 | lsrB | 1.316268 | 0.044154 | Y | -0.28482 | 0.621545 |
| P0A9L3 | fklB | 1.175144 | 0.045685 | Y | 0.246729 | 0.63484 |
| P0ABS1 | dksA | 1.788491 | 0.046564 | Y | 0.332091 | 0.673989 |
| P32680 | yjaG | 1.431228 | 0.02632 | Y | -0.21908 | 0.690318 |
| P0A725 | lpxC | 1.22745 | 0.004099 | Y | 0.129172 | 0.690472 |
| P0ADV7 | mlaC | 1.187597 | 0.033767 | Y | -0.18393 | 0.704139 |
| P0A927 | tsx | 2.005046 | 0.00036 | Y | -0.12972 | 0.718009 |
| P76403 | trhP | 1.060727 | 0.009909 | Y | -0.09164 | 0.780982 |
| P12008 | aroC | 1.288657 | 0.043098 | Y | -0.14855 | 0.790143 |
| P0A898 | ybeY | 1.831846 | 0.000433 | Y | -0.08242 | 0.806489 |
| P0AEI4 | rimO | 1.49232 | 0.008234 | Y | -0.10142 | 0.820471 |

| | | | | | | |
|---|---|---|---|---|---|---|
| P39180 | flu | 3.484661 | 1.41E-05 | Y | -0.08699 | 0.824205 |
| P04949 | fliC | 3.655465 | 0.001698 | Y | -0.12626 | 0.877122 |
| P33136 | mdoG | 1.130534 | 0.032001 | Y | -0.06102 | 0.892888 |
| P0AC81 | gloA | 1.783693 | 0.022495 | Y | -0.0858 | 0.896235 |
| P02924 | araF | 1.188544 | 0.02438 | Y | 0.049174 | 0.9123 |
| P76177 | ydgH | 1.66776 | 0.007046 | Y | -0.04853 | 0.920159 |
| P0AC51 | zur | 1.531258 | 0.003454 | Y | 0.035819 | 0.926983 |
| P0ACI0 | rob | 1.247464 | 0.042635 | Y | 0.045186 | 0.93303 |
| P39406 | rsmC | 1.235008 | 0.041791 | Y | 0.043731 | 0.934184 |
| P21367 | ycaC | 1.250443 | 0.000623 | Y | -0.01996 | 0.934293 |
| P0A850 | tig | 1.175986 | 0.018689 | Y | 0.031368 | 0.939825 |
| P02925 | rbsB | 1.581056 | 0.042357 | Y | -0.04642 | 0.945297 |
| P0A9X9 | cspA | 1.678779 | 0.000779 | Y | -0.0212 | 0.949743 |
| P0A8H6 | yihI | 1.01319 | 0.022109 | Y | -0.01972 | 0.957736 |
| P0AEE5 | mglB | 1.246589 | 0.049897 | Y | -0.0249 | 0.964384 |
| P42593 | fadH | 2.465445 | 6.30E-05 | Y | 0.003997 | 0.990728 |

**Table S2. Known CFX binding partners enriched and competed in chemoproteomics.** Known **CFX**-interacting proteins enriched and competed, but not meeting the definition for target proteins as statistically-significant hits in both conditions (n = 3). *AcrB is also present in Table S1.

| | | **5** vs DMSO | | (**5**+**CFX**) vs **5** | |
|---|---|---|---|---|---|
| Protein.ID | Gene | $log_2$(fold-change) | p-value | $log_2$(fold-change) | p-value |
| P0AES6 | gyrB | 0.302736 | 0.057871 | -0.36398 | 0.028592 |
| P31224 | acrB* | 1.009884 | 0.013468 | -0.503 | 0.156856 |
| P02930 | tolC | 0.828125 | 0.118241 | -0.27892 | 0.57298 |
| P06996 | ompC | 0.231276 | 0.616423 | -0.22585 | 0.624593 |

**Table S3. Full list of all identified proteins (separate Excel file).** Raw data are available via ProteomeXchange with identifier PXD068697

### Biological Methods

#### Bacterial strains, culture conditions and compound treatment

Bacterial strains, outlined in Table 1, were revived from a frozen stock, as an overnight culture grown on non-selective Muller Hinton Agar (MHA, Sigma-Aldrich, UK) supplemented with associated antibiotics at 37 °C for 18 h. Overnight cultures were grown in Muller Hinton Broth (MHB, Sigma-Aldrich, UK) except where stated, supplemented with associated antibiotics to late stationary phase at 37 °C and 200 RPM in a shaking incubator (Inova 42, New Brunswick Scientific, USA). Plates were incubated at 37 °C and 200 RPM in a shaking incubator with a BreatheEasy seal (Diversified Biotech, USA). Compounds were prepared as DMSO stocks for biological experiments except **CFX** which was dissolved in 10 mM HCl. All compounds were stored at -20 °C and thawed on the day of use.

**Table S4. Bacterial strains used in this work**

| **Bacterial strain** | **Description** | **Resistance Markers (Concentration)** | **Source** |
|---|---|---|---|
| *E. coli* MG1655 K12 | | | DSMZ (Germany) |
| *E. coli* MG1655 K12 *S83L-gyrA* | *E. coli* containing **CFX**-resistance mutation | | Orritt *et al.* 2022.[1] |
| *E. coli* MG1655 K12 *psulA-gfp* | *E. coli* containing GFP SOS reporter | Chloramphenicol (25 μg/mL) | Cheng *et al.* 2023[2] |
| *E. coli* BW25113 K12 Δ*yjdN* | *E. coli* single gene knockout of *yjdN.* | Kanamycin (50 μg/mL) | Baba *et al.* 2006[3] |
| *E. coli* BW25113 K12 Δ*ybaA* | *E. coli* single gene knockout of *ybaA.* | Kanamycin (50 μg/mL) | Baba *et al.* 2006[3] |
| *E. coli* BW25113 K12 Δ*pdxI* | *E. coli* single gene knockout of *pdxI.* | Kanamycin (50 μg/mL) | Baba *et al.* 2006[3] |
| *E. coli* BW25113 K12 Δ*elbB* | *E. coli* single gene knockout of *elbB.* | Kanamycin (50 μg/mL) | Baba *et al.* 2006[3] |

| | | | |
|---|---|---|---|
| T7 express *E. coli* | Chemically competent BL21 derived *E. coli* | | NEB (USA) |

**Construction of SOS reporter strains**

The *PsulA-gfp* plasmid was isolated from *E. coli* MG1655 K12 *PsulA-gfp* using a Monarch plasmid miniprep kit (NEB, USA) as per the manufacturer's instructions. *E. coli* K12 strains to be made competent were grown overnight in Lenox Broth (LB, Sigma-Aldrich, UK), and 50 µL subsequently subcultured to ~0.4 $OD_{600}$ in LB (10 mL). Cells were pelleted (5,000 × g, 5 min, 4 °C) and resuspended in 5 mL of Transformation and Storage Solution (TSS, 60 mM $CaCl_2$, 15% glycerol, 10 mM HEPES, pH 7.0, autoclaved) and incubated on ice (2 h). Cells were pelleted (5,000 × g, 5 min, 4 °C) and resuspended in 200 µL of TSS. 100 µL competent cells were then incubated with 5 µL of eluted *PsulA-gfp* plasmid on ice (30 min), heat shocked (42 °C, 30 s) and incubated in LB (37 °C, 1 h). Colonies were selected on LB Agar (LBA, Sigma-Aldrich, UK) with appropriate selection markers.

**Software analysis and plate reading**

$OD_{600}$ and fluorescence intensity readings were recorded in a CLARIOstar Plus microplate reader (BMG, UK). Measurements were background corrected against a non-inoculum control of MHB and normalised to DMSO control. GraphPad Prism 10.0.0 (Dotmatics, USA) was used to generate log dose-response curves and calculate SEM. Imaged gels were analysed in Empiria Studio software (LI-COR, USA). Surface Plasmon Resonance data was analysed using TraceDrawer (Ridgeview Instruments, Sweden).

**Minimum inhibitory concentration (MIC) assay**

MIC broth microdilution was performed according to CLSI methods M07-A11[4] in 384-well plates (Greiner Bio-One, UK). Two-fold serial dilutions of compounds in triplicate were prepared in MHB with a final volume of 25 µL and a no-compound growth control. A direct colony suspension was made by dispersing singular, well-isolated bacterial colonies from overnight revive plates in 3 mL sterile Phosphate-Buffered Saline (PBS) to achieve a turbidity of 0.5 McFarland standard (Oxoid, UK), approximately $10^8$ CFU/mL. The inoculum was vortexed and then further diluted 1:100 in MHB to achieve a final inoculum of $10^6$ CFU/mL. Inoculum (25 µL) was added to each well to achieve a final CFU/mL of $5 \times 10^5$, excluding no-inoculum sterility control which had only MHB added (final volume 50 µL, 1% (*v/v*) DMSO). Plates were incubated overnight for 18 h after which the background corrected $OD_{600}$ was measured. The MIC was recorded as the minimum concentration with mean normalised $OD_{600} < 0.1$.

#### SOS reporter assay

SOS response activation was measured in 384-well plates (Greiner Bio-One, UK) using SOS reporter strains. Two-fold serial dilutions of compounds in triplicate were prepared in MHB with a final volume of 25 µL and a no-compound growth control. Overnight cultures of *E. coli* maintaining *psulA-GFP* were diluted 8-fold and 25 µL added to each well (excluding no-inoculum control) to achieve $4 \times 10^7$ CFU (final volume 50 µL, 1% (*v/v*) DMSO). Plates were incubated for 6 h after which GFP fluorescence (Ex 375, Em 425) and $OD_{600}$ were measured, and background corrected $GFP/OD_{600}$ reported.

#### Probe 5 photoactivation

Probe **5** was diluted to 50 µM in 250 µl PBS and irradiated on ice in a UV Irradiator (Wavey Technologies, UK). Samples were kept on ice and in the dark when not irradiated. Samples were then analysed by High Performance Liquid Chromatography (HPLC) using an SPD-20A UV detector (Shimadzu, Japan) set to 280 nm and an ACE Equivalence 3, C18, 150 × 4.6 mm column (Avantor, USA), with a sample injection volume of 25 µL.

#### Affinity-based protein profiling (AfBPP)

Overnight cultures (WT or knockout mutants) were pelleted (35,000 × g, 10 min, 4 °C), washed twice in PBS, and resuspended in PBS to $OD_{600}$ of ~6, measured on a

Nanophotometer NP80. 1 mL aliquots of resuspension were incubated (500 rpm, 1 h, 37 °C) with required compound(s) or DMSO. Samples were UV-irradiated on ice for 12 min, where stated. Samples were centrifuged (17,000 × g, 5 min, 4 °C), the supernatant discarded, and pellets frozen at -20 °C. Pellets were resuspended in 300 µL lysis buffer (1% (v/v) Triton X-100, 1% (*w/v*) sodium dodecyl sulfate (SDS), EDTA-free complete protease inhibitor cocktail (1×, Roche) in PBS) and incubated on ice for 55 min. Samples were probe sonicated (40% amplitude, 2 min (10 s pulse, 10 s rest), Vibra-cell, Sonics, UK) on ice, then centrifuged (17,000 × g, 5 mins, 4 °C). The supernatant was removed and protein concentration was determined using the DC Protein Assay (Bio-Rad) as per the manufacturer's instructions. Samples were adjusted to 1 mg/mL in PBS (100 µL).

**CuACC ligation**

AzTB (10 mM, 1 µL), $CuSO_4$ (50 mM, 2 µL), TBTA (10 mM, 1 µL), TCEP (50 mM, 2 µL) were premixed, and added to 100 µL lysate. Reaction mixtures were incubated (500 rpm, 1 hr, RT), then quenched (5 mM EDTA) on ice. Proteins were precipitated by sequential addition of $H_2O$ (100 µL), MeOH (200 µL) and $CHCl_3$ (50 µL) to each sample at RT. Samples were vortexed and centrifuged (17,000 × g, 5 min, 4 °C). The top layer (MeOH/$H_2O$) was removed and 300 µL MeOH added, the samples were sonicated (5 min), centrifuged (17,000 × g, 5 min, 4 °C) and the resulting protein pellet washed with MeOH (300 µL × 3). Pellets were air dried (5 min), resuspended in 20 µL of 1% SDS in PBS and probe sonicated (10% amplitude, 10 s) on ice, then diluted to 1 mg/mL by adding 80 µL of PBS.

**In-gel fluorescence analysis**

SDS-PAGE samples were prepared by sequential addition of 5 µL 4 × NuPAGE LDS Sample Buffer (Thermofisher, UK), 0.5 M DTT (2 µL) and 13 µL of sample. Samples were heated for 5 min at 95 °C, loaded onto 12% Bis-Tris gels and separated by electrophoresis using an Invitrogen XCell SureLock Electrophoresis Mini-Cell Tank with an Invitrogen PowerEase Touch 350W Power Supply (Thermofisher, UK). TAMRA fluorescence was imaged at 520 nm on an Odyssey M imager (LI-COR). Total protein was imaged after staining with QuickBlue Protein Stain (Stratech Scientific Ltd, UK) at 700 nm on an Odyssey M imager.

#### Pull-down, on bead digest and LC-MS/MS analysis

CuAAC ligation and protein precipitation was performed as described previously with the reaction scaled up to 1 mg of total protein for each condition and protein concentrations adjusted to 2.5 mg/mL. 100 µL of 50% slurry NeutrAvidin Agarose Resin (Thermo Scientific, UK) per 1 mg of protein was washed three times with excess 0.2% SDS in PBS and isolated by centrifugation (3,000 × g, 2 mins, 4 °C). Beads were aliquoted and incubated with samples with shaking (1,200 rpm, 2 h, RT). Samples were centrifuged, then the supernatant removed and beads washed three times as before. Beads were washed in Urea-AmBic buffer (8 M Urea in 100 mM ammonium bicarbonate, pH 7.8, 2 × 100 µL), resuspended in 1.5 × volume Urea-AmBic buffer and incubated with shaking (700 rpm, 10 min, RT). TCEP was added (10 mM) and samples incubated (30 min, RT), then 2-chloroacetamide was added (50 mM) and samples incubated (30 min, RT). Bound proteins were pre-digested using 1 µg LysC per 100 µg protein (700 rpm, 2 h, 37 °C). Urea was diluted to 2 M in 100 mM AmBic, $CaCl_2$ was added (2 mM), and 1 µg trypsin per 40 µg protein was added before incubation (700 rpm, 16 h, 37 °C), then 5 % formic acid was added to quench the reaction. Samples were centrifuged (17000 × g, 30 min, 4 °C), the supernatant desalted on C18 columns, then bound peptides eluted with 50% acetonitrile + 0.1% TFA. Peptides were dried and analysed on a Q Exactive Hybrid Quadrupole-Orbitrap (Thermofisher, UK) for liquid chromatography with tandem mass spectrometry (LC-MS/MS).

#### Chemoproteomics data analysis

Raw files were searched using FragPipe (v.22.0) against the *E. coli* K12 proteome (UP000000625, downloaded on 06.04.2024 from UniProt) using the built in LFQ-MBR workflow.[4] All standard parameters were kept, and ‘stricttrypsin’ with up to two missed cleavages was applied for *in silico* digestion. Methionine oxidation and N-terminal acetylation were specified as variable modifications, while cysteine carbamidomethylation was set as a fixed modification. Protein level output data were further analysed using R (v.4.3.1, R https://www.R-project.org/). Proteins that were detected at less than 30% across all samples were removed. MaxLFQ intensities were $log_2$ transformed and missing values imputed using the MinDet function. Statistical analysis was performed using limma.[5]

### Purified protein production

pET28a plasmids were synthesised by GenScript (GenScript Biotech, USA), containing *yjdn*, *ybaA, ybaA*-E40A, or *ybaA*-E40K with a C-terminal $His_{10}$-tag, and transformed into T7 express *E. coli* (NEB, USA). For expression, overnight cultures were diluted 1:50 in LB (BD Difco, USA) supplemented with kanamycin (50 µg/mL) and cultured to an $OD_{600}$ of 0.7, then induced by addition of IPTG to a final concentration of 1 mM, and cells grown overnight at 18 °C. Cells were harvested by centrifugation (5,000 × g, 30 min, 4 °C) and pellets resuspended in lysis buffer (50 mM Tris pH 8.0, 300 mM NaCl). Cells were lysed by three passes through an Emulsiflex-C3 cell disruptor (Avestin, Canada), then cell debris removed by centrifugation (35,000 × g, 30 min, 4 °C). The supernatant was incubated with Ni-NTA resin (GE Healthcare, USA) (2 h, 4 °C), which was sequentially washed with Ni Wash Buffer (4x50 mL, 50 mM Tris pH 8.0, 300 mM NaCl, 40 mM imidazole). Bound protein was then eluted with Ni Elution Buffer (50 mM Tris pH 8.0, 300 mM NaCl, 250 mM imidazole). Protein-containing fractions were pooled and concentrated using an Amicon Ultra centrifugal filter (Merck, USA) (10 kDa cut-off) and further purified by size exclusion chromatography on a Superdex 75 HiLoad 16/600 gel filtration column (GE healthcare, USA). Peak fractions were pooled, concentrated and either immediately used or flash-frozen in liquid nitrogen for storage at -80 °C.

### *In vitro* crosslinking with purified proteins

Purified protein (5 µM) was incubated with probe **5** (5 µM) with or without **CFX** (100 µM) or **6** (100 µM) competition in reaction buffer (20 mM Tris,150 mM NaCl, 100 µL total volume). Samples were incubated for 1 h, 37 °C, 500 rpm, then UV-irradiated on ice for 12 min where stated. AzTB (10 mM, 1 µL), $CuSO_4$ (50 mM, 2 µL), TBTA (10 mM, 1 µL), TCEP (50 mM, 2 µL) were premixed and added to 100 µL sample. Reaction mixtures were incubated (500 rpm, 1 h, RT), then quenched (5 mM EDTA) on ice. Samples were visualised by in-gel fluorescence as described for AfBPP in live *E. coli*.

**Surface plasmon resonance (SPR)**

SPR experiments were conducted on a Biacore T200 (Cytiva, USA) at 37 °C with a flow rate of 10 µL/min. The running buffer consisted of 20 mM Tris and 150 mM NaCl, pH 7.4, filtered and degassed prior to use. A Sensor Chip NTA (Cytiva, USA) was conditioned by injecting 350 mM EDTA in running buffer for 120 s at a flow rate of 10 µL/min. The chip surface was saturated with nickel by injecting 0.5 mM $NiCl_2$ at a flow rate of 10 µL/min for 60 s, and then with 3 mM EDTA at a flow rate of 10 µL/min for 30 s. $His_{10}$-tagged proteins were captured on the nickel-saturated surface by injecting solutions at a concentration of 0.01 µg/mL. The injection was performed for 60 s at a flow rate of 10 µL/min. A dose-response of **CFX** was injected over the captured protein at a flow rate of 10 µL/min for 120 s.

***In vitro* crosslinking and LC-MS/MS binding site identification**

Purified protein (15 µg) was incubated with or without probe **5** (5 µM) in reaction buffer (20 mM Tris,150 mM NaCl, 300 µL total volume). Samples were incubated for 1 h, 37 °C, 500 rpm then UV-irradiated on ice for 12 min. Samples were diluted in 8 M urea in 100 mM AmBic buffer, pH 7.8, 2 × 100 µL) and incubated (650 rpm, 10 min, RT). TCEP was added (10 mM) and samples incubated (650 rpm, 30 min, RT), then 2-chloroacetamide was added (50 mM) and samples incubated (650 rpm, 30 min, RT). Samples were pre-digested using 1 µg LysC per 100 µg protein (650 rpm, 2 h, 37 °C). Urea was diluted to 2 M in 100 mM AmBic, $CaCl_2$ was added (2 mM), and 1 µg trypsin per 40 µg protein was added before incubation (800 rpm, 18 h, 37 °C), then 5 % formic acid was added to quench the reaction. Samples were centrifuged (17,000 × g, 30 min, 4 °C), and the supernatant desalted using C18 StageTips, then eluted with 50% acetonitrile 0.1% TFA. Peptides were dried and analysed on a Q Exactive Hybrid Quadrupole-Orbitrap LC-MS/MS. Data was processed using Proteome Discoverer

(ThermoFisher, UK), with the carbene added as a variable modification (mass = 423.195820), and a search was conducted on all residues except cysteine.

#### *In silico* docking

**CFX** was docked into the *S. flexneri* YbaA structure (PDB: 2okq) using Flare™ V10.0.1 (Cresset, UK), defining the protonation state via full preparation on the protein, with intelligent capping.

### Chemical Methods

#### General Information

Materials were purchased from commercial suppliers and used as received. Analytical thin-layer chromatography (TLC) was performed on 0.25 mm silica gel 60 F254 pre-coated plates 0.25 mm (Merck, UK) and visualized under ultraviolet light (254 and 365 nm). Purification by column chromatography was carried out using a CombiFlash R*f* automated column system with RediSep silver disposable flash columns (Teledyne, USA).

$^1$H, $^{13}$C, and $^{19}$F NMR spectra were recorded at room temperature at 400 MHz, 101 MHz, and 376 MHz, respectively (Bruker, USA). Chemical shifts are reported as parts per million (δ) using trimethylsilane (TMS) and the peak of the residual solvent proton signals as internal reference. Coupling constants (*J*) are reported in hertz (Hz) and averaged for interacting protons. Low-resolution mass spectroscopy (LRMS) and high-resolution mass spectroscopy (HRMS) was recorded on a BioAccord (Waters, USA). HPLC analysis was conducted using an SPD-20A UV detector (Shimadzu) with 280 nm detection, and an ACE Equivalence 3, C18, 150 × 4.6 mm column (Avantor).

#### Synthetic Procedures

***Ethyl 1-(2-(3-(but-3-yn-1-yl)-3H-diazirin-3-yl)ethyl)-6,7-difluoro-4-oxo-1,4-dihydroquinoline-3-carboxylate*** (**2c**)

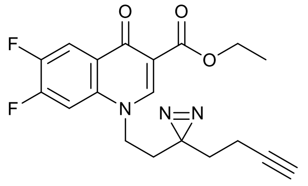

Ethyl (*Z*)-2-(2-amino-4,5-difluorobenzoyl)-3-ethoxyacrylate (**2a**) and ethyl 6,7-difluoro-4-oxo-1,4-dihydroquinoline-3-carboxylate (**2b**) were prepared as previously described.[6] **2b** (100 mg, 0.39 mmol) and $K_2CO_3$ (55 mg, 0.39 mmol) were suspended in anhydrous DMF (2 mL) and 3-(but-3-yn-1-yl)-3-(2-iodoethyl)-3H-diazirine (98 mg, 0.39 mmol) was added. The reaction was stirred at RT for 16 h after which sat. aq. $NH_4Cl$ (20 mL) was added, and the suspension was stirred for 10 min. The product was extracted with DCM (3 × 10 mL), dried ($MgSO_4$), and purified by flash column chromatography, 0–60% EtOAc in petroleum ether to give a white solid (36 mg, 76%). $R_f$ = 0.85 ($SiO_2$; DCM:MeOH, 90:10); **$^1$H NMR** (400 MHz, $CDCl_3$) δ 8.53 (s, 1H), 8.32 (dd, *J* = 10.4, 8.8 Hz, 1H), 7.15 (dd, *J* = 11.0, 6.1 Hz, 1H), 4.41 (q, *J* = 7.1 Hz, 2H), 4.04–3.95 (m, 2H), 2.11–2.03 (m, 5H), 1.61 (m, 2H), 1.42 (t, *J* = 7.1 Hz, 3H); **$^{13}$C NMR** (101 MHz, $CDCl_3$) δ 172.5, 165.1, 153.5 (dd, *J* = 258.6, 15.2 Hz), 149.1, 148.6 (dd, *J* = 218.2, 13.1 Hz), 135.3, 126.5, 116.2 (d, *J* = 18.6 Hz), 111.5, 104.3 (d, *J* = 22.6 Hz), 82.4, 70.1, 61.2, 48.8, 32.5, 32.2, 25.9, 14.4, 13.2; **$^{19}$F NMR** (376 MHz, $CDCl_3$) δ: -126.86, -138.18; **LRMS** m/z ($ESI^+$) 374 ($[M+H]^+$); **HRMS** m/z ($ESI^+$) found 374.1307 $C_{19}H_{18}F_2N_3O_3$ ($[M+H]^+$) requires 374.1311.

***1-(2-(3-(But-3-yn-1-yl)-3H-diazirin-3-yl)ethyl)-6-fluoro-4-oxo-7-(piperazin-1-yl)-1,4-dihydroquinoline-3-carboxylic acid*** (**2**)

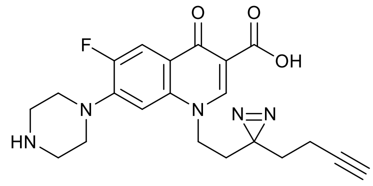

Ethyl 1-(2-(3-(but-3-yn-1-yl)-3H-diazirin-3-yl)ethyl)-6,7-difluoro-4-oxo-1,4-dihydroquinoline-3-carboxylate (**2c**) (70 mg, 0.19 mmol) and piperazine (65 mg, 0.75 mmol) were stirred in anhydrous MeCN (3 mL) at 50 °C. After 36 h, aq. 1 M NaOH (2 mL) and $H_2O$ (5 mL) was added and stirred for 2 h, after which the solution reacidified to pH 7 (aq. 1M HCl). MeCN was removed *in vacuo* and the product precipitated at 4 °C. The solid was filtered and washed with $H_2O$ (10 mL) and $Et_2O$ (10 mL) to give a white solid (48 mg, 62%). $R_f$ = 0.23 ($SiO_2$; DCM:MeOH, 80:20). **$^1$H NMR** (400 MHz, $CDCl_3$) δ 8.72 (s, 1H), 8.05 (d, *J* = 13.1 Hz, 1H), 6.61 (d, *J* = 6.8 Hz, 1H), 4.19 (t, *J* = 7.9 Hz, 2H), 3.29 (t, *J* = 4.8 Hz, 4H), 3.11 (t, *J* = 4.8 Hz, 4H), 2.08 (t, *J* = 5.6 Hz, 3H), 2.04 (d, *J* = 8.0 Hz, 2H), 1.65 (t, *J* = 6.8 Hz, 2H); **$^{13}$C NMR** (151 MHz, $CDCl_3$:MeOD) δ 177.1 (d, *J* = 2.7 Hz), 167.6, 153.6 (d, *J* = 252.2 Hz), 148.0, 146.5 (d, *J* = 10.2 Hz), 137.0, 120.5 (d, *J* = 8.0 Hz), 112.9 (d, *J* = 23.4 Hz), 108.2, 103.6 (d, *J* = 3.6 Hz), 82.3,

70.1, 50.5 (d, *J* = 5.0 Hz), 49.0, 45.4, 32.8, 31.7, 29.7, 26.0, 13.1; **$^{19}$F NMR** (376 MHz, $CDCl_3$) δ: -120.18; **LRMS** m/z (ESI$^+$) 412 ([M+H]$^+$); **HRMS** m/z (ESI$^+$) found 412.1794, $C_{21}H_{23}FN_5O_3$ ([M+H]$^+$) requires 412.1779; **HPLC** Retention time 7.6 min, 95% (280 nm).

***Ethyl 6,7-difluoro-4-oxo-1-(4-(3-(trifluoromethyl)-3H-diazirin-3-yl)benzyl)-1,4-dihydroquinoline-3-carboxylate*** (**3a**)

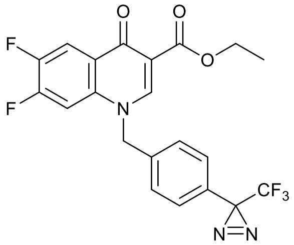

Ethyl 6,7-difluoro-4-oxo-1,4-dihydroquinoline-3-carboxylate (**2b**) (100 mg, 0.39 mmol) and $K_2CO_3$ (109 mg, 0.79 mmol) were suspended in anhydrous DMF (2 mL) and 4-[3-(trifluoromethyl)-3H-diazirin-3-yl]benzyl bromide (110 mg, 0.39 mmol) was added. The reaction was stirred at RT for 16 h after which sat. aq. $NH_4Cl$. (20 mL) was added, and the suspension was stirred for 10 min. The product was extracted with DCM (3 × 10 mL), dried ($MgSO_4$), and purified by flash column chromatography, 0–5% MeOH in DCM to give a white solid (136 mg, 76%). $R_f$ = 0.63 ($SiO_2$; DCM:MeOH, 95:5); **$^1$H NMR** (400 MHz, $CDCl_3$) δ 8.55 (s, 1H), 8.27 (dd, *J* = 10.3, 8.7 Hz, 1H), 7.26–7.15 (m, 4H), 7.02 (dd, *J* = 11.0, 6.1 Hz, 1H), 5.37 (s, 2H), 4.39 (q, *J* = 7.1 Hz, 2H), 1.40 (t, *J* = 7.1 Hz, 3H); **$^{13}$C NMR** (101 MHz, $CDCl_3$) δ 172.6, 165.1, 153.4 (dd, *J* = 258.6, 15.2), 149.8, 148.8 (dd, *J* = 257.6, 14.1), 135.9 (d, *J* = 7.9 Hz), 135.2, 130.1, 127.7 (2C), 126.5 (2C), 126.4, 121.9 (q, *J* = 275.0 Hz), 115.8 (d, *J* = 20.6 Hz), 111.5, 105.4 (d, *J* = 22.6 Hz), 77.2, 61.3, 57.2, 28.4 (q, *J* = 40.8 Hz), 14.4; **$^{19}$F NMR** (376 MHz, $CDCl_3$) δ: -65.16 -126.47, -138.00. **LRMS** m/z (ESI$^+$) 452 ([M+H]$^+$); **HRMS** m/z (ESI$^+$) found 452.1042, $C_{21}H_{15}F_5N_3O_3$ ([M+H]$^+$) requires 452.1028.

***6-Fluoro-4-oxo-7-(piperazin-1-yl)-1-(4-(3-(trifluoromethyl)-3H-diazirin-3-yl)benzyl)-1,4-dihydroquinoline-3-carboxylic acid*** (**3b**)

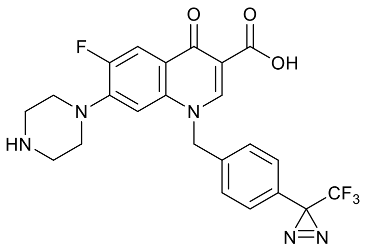

Ethyl 6,7-difluoro-4-oxo-1-(4-(3-(trifluoromethyl)-3H-diazirin-3-yl)benzyl)-1,4-dihydroquinoline-3-carboxylate (**3a**) (109 mg, 0.24 mmol) and piperazine (83 mg, 0.96 mmol) were stirred in anhydrous MeCN (5 mL) at 40 °C. After 36 h, aq. 1M NaOH (2 mL) and $H_2O$ (5 mL) was added and stirred for a further 3 h after which the solution neutralised to pH 7 (aq. 1 M HCl). MeCN was removed *in vacuo* and the product was left to precipitate in a 4 °C fridge. The solid was filtered and washed with $H_2O$ (10 mL) and $Et_2O$ (10 mL) to give a white solid (93 mg, 78%). $R_f$ = 0.28 ($SiO_2$; DCM:MeOH, 80:20). **$^1$H NMR** (400 MHz, acetic-*d₄*) δ 9.24 (s, 1H), 8.09 (d, *J* = 12.7 Hz, 1H), 7.50 (d, *J* = 8.0 Hz, 2H), 7.31 (d, *J* = 8.0 Hz, 2H), 7.10 (d, *J* = 6.8 Hz, 1H), 5.83 (s, 2H), 3.52 (s, 8H); **$^{13}$C NMR** (101 MHz, acetic-$d_4$) δ 168.9, 160.3, 153.5 (d, *J* = 243.2 Hz), 150.2, 144.8 (d, *J* = 11.1 Hz), 137.7, 136.7, 129.2, 127.6 (2C), 127.2 (2C), 120.9 (q, *J* = 47.47 Hz), 123.4, 112.5, 112.3, 106.7, 57.4, 46.4 (2C), 43.3 (2C)[1]; **$^{19}$F NMR** (376 MHz, acetic-$d_4$) δ: -66.35 -122.18. **LRMS** m/z ($ESI^+$) 490 ($[M+H]^+$); **HRMS** m/z ($ESI^+$) found 490.1519, $C_{23}H_{20}F_4N_5O_3$ ($[M+H]^+$) requires 490.1497.

***6-Fluoro-4-oxo-7-(4-(prop-2-yn-1-yl)piperazin-1-yl)-1-(4-(3-(trifluoromethyl)-3H-diazirin-3-yl)benzyl)-1,4-dihydroquinoline-3-carboxylic acid*** (**3**)

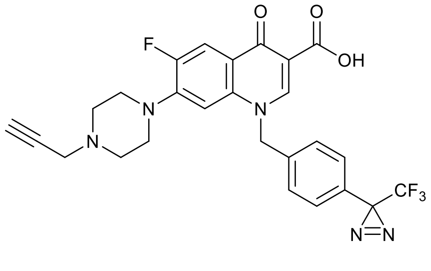

6-Fluoro-4-oxo-7-(piperazin-1-yl)-1-(4-(3-(trifluoromethyl)-3H-diazirin-3-yl)benzyl)-1,4-dihydroquinoline-3-carboxylic acid (**3b**) (40 mg, 0.082 mmol) and $K_2CO_3$ (17 mg,

[1] No signal observed for C-diazirine, ~25 ppm, due to fluorine coupling.

0.12 mmol) were suspended in anhydrous MeCN (2 mL) and propargyl bromide (15 mg, 0.12 mmol) was added. The reaction was stirred at RT for 16 h after which aq. sat. $NH_4Cl$ (10 mL) was added, and the suspension was stirred for 10 min. The product was extracted with DCM (3 × 5 mL), dried ($MgSO_4$), and purified by flash column chromatography, 0–5% MeOH in DCM and the product washed with $Et_2O$ to afford a white solid (23 mg, 53%). $R_f$ = 0.42 ($SiO_2$; DCM:MeOH, 19:1). **$^1$H NMR** (400 MHz, $CDCl_3$) δ 14.96 (s, 1H), 8.80 (s, 1H), 8.03 (d, *J* = 13.1 Hz, 1H), 7.22 (s, 4H), 6.60 (d, *J* = 6.9 Hz, 1H), 5.46 (s, 2H), 3.36 (d, *J* = 2.5 Hz, 2H), 3.17 (t, *J* = 4.7, 4H), 2.70 (t, *J* = 4.7, 4H), 2.30 (t, *J* = 2.5 Hz, 1H);**$^{13}$C NMR** (101 MHz, $CDCl_3$) δ 177.3, 167.0, 153.5 (d, *J* = 252.3 Hz), 148.4, 145.8 (d, *J* = 11.11 Hz), 137.4, 135.3, 130.3, 127.7 (2C), 126.7 (2C), 120.6 (q, *J* = 5.05 Hz), 112.8 (d, *J* = 23.2 Hz), 108.6, 104.9, 78.1, 73.7, 58.0, 51.3 (2C), 49.5 (d, *J* = 5.2 Hz, 2C), 46.8[2]; **$^{19}$F NMR** (376 MHz, $CDCl_3$) δ: -65.12, -120.06; **LRMS** m/z ($ESI^+$) 528 ($[M+H]^+$); **HRMS** m/z ($ESI^+$) found 528.1651, $C_{26}H_{22}F_4N_5O_3$ ($[M+H]^+$) requires 528.1653; **HPLC** Retention time 9.9 min, 97% (280 nm).

***Ethyl 6,7-difluoro-1-(2-methylbut-3-yn-2-yl)-4-oxo-1,4-dihydroquinoline-3-carboxylate*** (**4b**)

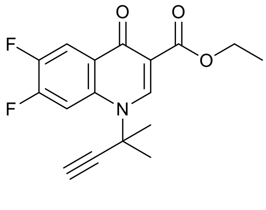

Ethyl (*Z*)-3-(dimethylamino)-2-(2,4,5-trifluorobenzoyl)acrylate (**4a**) was prepared as previously described.[6]

*Step 1:* **4a** (880 mg, 2.9 mmol) was dissolved in EtOH (5 mL) and $Et_2O$ in (5 mL) and 2-methyl-3-butyn-2-amine (490 mg, 5.80 mmol) was added. The reaction was stirred at RT for 4 h after the solvent was removed *in vacuo* to give the crude intermediate as a yellow oil. $R_f$ = 0.80 ($SiO_2$; PetEth:EtOAc, 1:1)

---

[2] No signal observed for C-diazirine, ~25 ppm, assumed due to fluorine coupling.

*Step 2:* The crude intermediate was dissolved in DMF (10 mL) and $K_2CO_3$ (1.6 g, 11.7 mmol) added. The reaction was stirred at 100 °C for 4 h after which sat. aq. $NH_4Cl$ (50 mL) was added and the product extracted with DCM (3 × 20 mL), dried ($MgSO_4$), and purified by flash column chromatography, 25–100% EtOAc in petroleum ether to give a yellow solid (450 mg, 48%). $R_f$ = 0.24 ($SiO_2$; PetEth:EtOAc, 1:1); **$^1$H NMR** (400 MHz, $CDCl_3$) δ 8.82 (s, 1H), 8.39–8.28 (m, 2H), 4.41 (q, *J* = 7.1 Hz, 2H), 2.84 (s, 1H), 2.09 (s, 6H), 1.42 (t, *J* = 7.1 Hz, 3H); **$^{13}$C NMR** (101 MHz, $CDCl_3$) δ 172.3, 165.9, 152.0 (dd, *J* = 253.8, 14.8 Hz), 148.3 (dd, *J* = 251.8, 13.8 Hz), 144.7, 135.2, 127.7 (d, *J* = 6.0 Hz), 115.6 (dd, *J* = 18.4, 2.5 Hz), 110.5, 108.9 (d, *J* = 24.3 Hz), 83.5, 76.8, 61.2, 58.0, 30.5, 14.4; **$^{19}$F NMR** (376 MHz, $CDCl_3$) δ: -127.88, -138.74; **LRMS** m/z (ESI$^+$) 320 ([M+H]$^+$); **HRMS** m/z (ESI$^+$) found 661.1913 $C_{34}H_{30}F_4N_2O_6Na$ ([2M+Na]$^+$) requires 661.1932.

***6-Fluoro-1-(2-methylbut-3-yn-2-yl)-4-oxo-7-(piperazin-1-yl)-1,4-dihydroquinoline-3-carboxylic acid*** (**4b**)

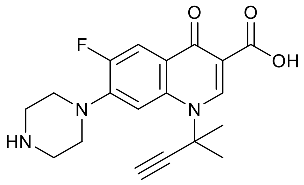

Ethyl 6,7-difluoro-1-(2-methylbut-3-yn-2-yl)-4-oxo-1,4-dihydroquinoline-3-carboxylate (**4b**) (200 mg, 0.62 mmol) and piperazine (220 mg, 0.25 mmol) were stirred in anhydrous MeCN (10 mL) at 50 °C. After 24 h, aq. 1M NaOH (5 mL) and $H_2O$ (5 mL) were added and stirred for 2 h, after which the solution reacidified to pH 7 (aq. 1M HCl). MeCN was removed *in vacuo* and the solid was filtered and washed with $H_2O$ (10 mL) and $Et_2O$ (10 mL) to give a pink solid, 200 mg (90%). $R_f$ = 0.10 ($SiO_2$; DCM:MeOH, 80:20); **$^1$H NMR** (400 MHz, acetic) δ 9.07 (s, 1H), 8.26 (d, *J* = 7.2 Hz, 1H), 8.14 (d, *J* = 12.9 Hz, 1H), 3.70 (dt, *J* = 6.6, 4.1 Hz, 4H), 3.63 (dt, *J* = 7.9, 4.2 Hz, 4H), 3.43 (s, 1H), 2.23 (s, 6H); **$^{13}$C NMR** (101 MHz, acetic) δ 177.3, 168.9, 153.3 (d, *J* = 250.7 Hz), 144.66, 143.41 (d, *J* = 10.5 Hz), 136.9, 122.2 (d, *J* = 7.7 Hz), 112.4 (d, *J* = 22.9 Hz), 110.2, 106.9, 83.7, 77.6, 59.6, 46.7, 43.4, 29.8; **$^{19}$F NMR** (376 MHz, acetic) δ: -122.52; **LRMS** m/z (ESI$^+$) 358 ([M+H]$^+$); **HRMS** m/z (ESI$^+$) found 385.1575,

$C_{19}H_{21}FN_3O_3$ ([M+H]$^+$) requires 358.1562; **HPLC** Retention time 7.03 min, 99% (280 nm).

***6-Fluoro-1-(2-methylbut-3-yn-2-yl)-4-oxo-7-(4-(4-(3-(trifluoromethyl)-3H-diazirin-3-yl)benzyl)piperazin-1-yl)-1,4-dihydroquinoline-3-carboxylic acid*** (**4**)

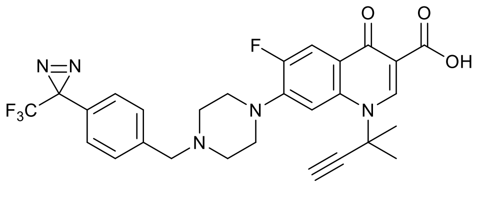

6-Fluoro-1-(2-methylbut-3-yn-2-yl)-4-oxo-7-(piperazin-1-yl)-1,4-dihydroquinoline-3-carboxylic acid (**4c**) (50 mg, 0.14 mmol) and $K_2CO_3$ (19 mg, 0.14 mmol) were suspended in anhydrous MeCN (2 mL) and 4-[3-(trifluoromethyl)-3H-diazirin-3-yl]benzyl bromide (59 mg, 0.21 mmol) was added. The reaction was stirred at RT for 24 h after which sat. aq. $NH_4Cl$ (10 mL) was added, and the suspension was stirred for 10 min. The product was extracted with DCM (3 × 5 mL), dried ($MgSO_4$), and purified by flash column chromatography, 0–5% MeOH in DCM to afford a white crystalline solid (50 mg, 64%). $R_f$ = 0.78 ($SiO_2$; DCM:MeOH, 9:1); **$^1$H NMR** (400 MHz, $CDCl_3$) δ 15.02 (s, 1H), 8.94 (s, 1H), 8.08 (d, *J* = 13.0 Hz, 1H), 7.95 (d, *J* = 7.1 Hz, 1H), 7.41 (d, *J* = 8.3 Hz, 1H), 7.18 (d, *J* = 7.8 Hz, 1H), 3.61 (s, 2H), 3.37–3.30 (m, 4H), 2.83 (s, 1H), 2.70–2.64 (m, 4H), 2.12 (s, 6H); **$^{13}$C NMR** (101 MHz, $CDCl_3$) δ 176.8, 167.4, 153.3 (d, *J* = 252.2 Hz), 144.6 (d, *J* = 10.1 Hz), 143.2, 139.7, 136.8, 129.5, 128.2, 126.6, 123.5, 121.7 (q, *J* = 6.7 Hz), 112.6 (d, *J* = 23.1 Hz), 109.0, 107.8, 83.7, 77.2, 76.6, 62.3, 58.6, 52.8, 49.9 (d, *J* = 4.9 Hz), 30.8; **$^{19}$F NMR** (376 MHz, $CDCl_3$) δ: -65.23, -120.85; **LRMS** m/z (ESI$^+$) 556 ([M+H]$^+$); **HRMS** m/z (ESI$^+$) found 556.1977, $C_{28}H_{26}F_4N_5O_3$ ([M+H]$^+$) requires 556.1966; **HPLC** Retention time 9.8 min, 98% (280 nm).

***7-(4-(2-(3-(But-3-yn-1-yl)-3H-diazirin-3-yl)ethyl)piperazin-1-yl)-1-cyclopropyl-6-fluoro-4-oxo-1,4-dihydroquinoline-3-carboxylic acid*** (**5**)

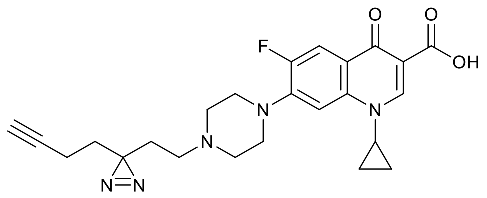

1-Cyclopropyl-6-fluoro-4-oxo-7-(piperazin-1-yl)-1,4-dihydroquinoline-3-carboxylic acid (**CFX**) (100 mg, 0.30 mmol) and $K_2CO_3$ (63 mg, 0.45 mmol) were suspended in anhydrous MeCN (2 mL) and 3-(but-3-yn-1-yl)-3-(2-iodoethyl)-3H-diazirine (75 mg, 0.30 mmol) was added. The reaction was stirred at 50 °C for 16 h after which sat. aq. $NH_4Cl$ (20 mL) was added, and the suspension was stirred for 10 min. The product was extracted with DCM (3 × 10 mL), dried ($MgSO_4$), and purified by flash column chromatography, 0–5% MeOH in DCM and the product washed with $Et_2O$ to afford a white solid (27 mg, 43%). $R_f$ = 0.60 ($SiO_2$; DCM:MeOH, 19:1). **$^1$H NMR** (400 MHz, $CDCl_3$) δ 15.00 (s, 1H), 8.74 (s, 1H), 7.97 (d, *J* = 13.1 Hz, 1H), 7.35 (d, *J* = 7.1 Hz, 1H), 3.55 (tt, *J* = 7.2, 4.0 Hz, 1H), 3.35 (t, *J* = 5.0 Hz, 4H), 2.64 (t, *J* = 5.0 Hz, 4H), 2.31 (t, *J* = 7.4 Hz, 2H), 2.09–1.96 (m, 3H), 1.80–1.61 (m, 3H), 1.42–1.37 (m, 2H), 1.29–1.16 (m, 2H); **$^{13}$C NMR** (101 MHz, $CDCl_3$) δ 177.1, 167.1, 153.7 (d, *J* = 251.5 Hz), 147.4, 145.9 (d, *J* = 10.1), 139.1, 119.8 (d, *J* = 7.8 Hz), 112.4 (d, *J* = 23.5 Hz), 108.1, 104.8 (d, *J* = 3.5 Hz), 82.8, 77.3, 69.2, 52.6, 52.6 (d, *J* = 9.4 Hz, 2C), 49.8 (d, *J* = 5.1 Hz, 2C), 35.3, 32.4, 30.3, 27.3, 13.3, 8.2; **$^{19}$F NMR** (376 MHz, $CDCl_3$) δ: -120.70; **LRMS** m/z (ESI$^+$) 452 ([M+H]$^+$); **HRMS** m/z (ESI$^+$) found 452.2104, $C_{24}H_{27}FN_5O_3$ ([M+H]$^+$) requires 452.2092; **HPLC** Retention time 8.3 min, 96% (280 nm).

**1-Cyclopropyl-7-(dimethylamino)-6-fluoro-4-oxo-1,4-dihydroquinoline-3-carboxylic acid (6)**

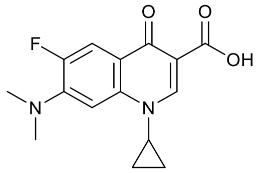

Ethyl 1-cyclopropyl-6,7-difluoro-4-oxo-1,4-dihydroquinoline-3-carboxylate (**6a**) was prepared as previously described.[6] **6a** (50 mg, 0.17 mmol) was dissolved in MeCN (1.7 mL), dimethylamine (33% in EtOH, 120 µL, 0.68 mmol) was added and the solution stirred at 80 ºC for 16 h. Aq. 1M NaOH (0.34 mL) was added and the solution stirred at 80 ºC for 2 h. The reaction mixture was diluted with 1 M HCl (0.51 mL) and extracted with DCM (3 × 5 mL), washed with brine, dried ($Na_2SO_4$), and purified by flash column chromatography with 0-5% MeOH in DCM (+0.1% FA) to afford the title compound as white solid (34 mg, 0.12 mmol, 68%). $R_f$ = 0.5 ($SiO_2$, DCM:MeOH, 19:1); **$^1$H NMR** (600 MHz, $CDCl_3$) δ 15.25 (s, 1H), 8.73 (s, 1H), 7.96 (d, *J* = 14.1 Hz, 1H), 7.12 (d, *J* = 7.5 Hz, 1H), 3.51 (tt, *J* = 7.1, 4.0 Hz, 1H), 3.16 (d, *J* = 1.9 Hz, 6H), 1.37 (dddd, *J* = 7.4, 6.4, 5.2, 0.9 Hz, 2H), 1.23 – 1.17 (m, 2H); **$^{13}$C NMR** (151 MHz, $CDCl_3$) δ 177.07 (d, *J* = 2.9 Hz), 167.59, 152.41 (d, *J* = 250.1 Hz), 147.31, 145.67 (d, *J* = 10.4 Hz), 139.56, 117.69 (d, *J* = 7.7 Hz), 112.59 (d, *J* = 24.1 Hz), 107.99, 102.02 (d, *J* = 4.5 Hz), 42.39 (d, *J* = 6.5 Hz), 35.31, 8.30; **$^{19}$F NMR** (377 MHz, $CDCl_3$) δ -121.80 (dd, *J* = 14.0, 7.3 Hz); **HRMS** *m/z* ($ESI^+$) found 291.1133, $C_{15}H_{16}FN_2O_3$ ($[M+H]^+$) requires 291.1139; **HPLC** Retention time 10.9 min, >99% (280 nm).

### NMR Spectra of Final Compounds

#### *1-(2-(3-(but-3-yn-1-yl)-3H-diazirin-3-yl)ethyl)-6-fluoro-4-oxo-7-(piperazin-1-yl)-1,4-dihydroquinoline-3-carboxylic acid* (**2**)

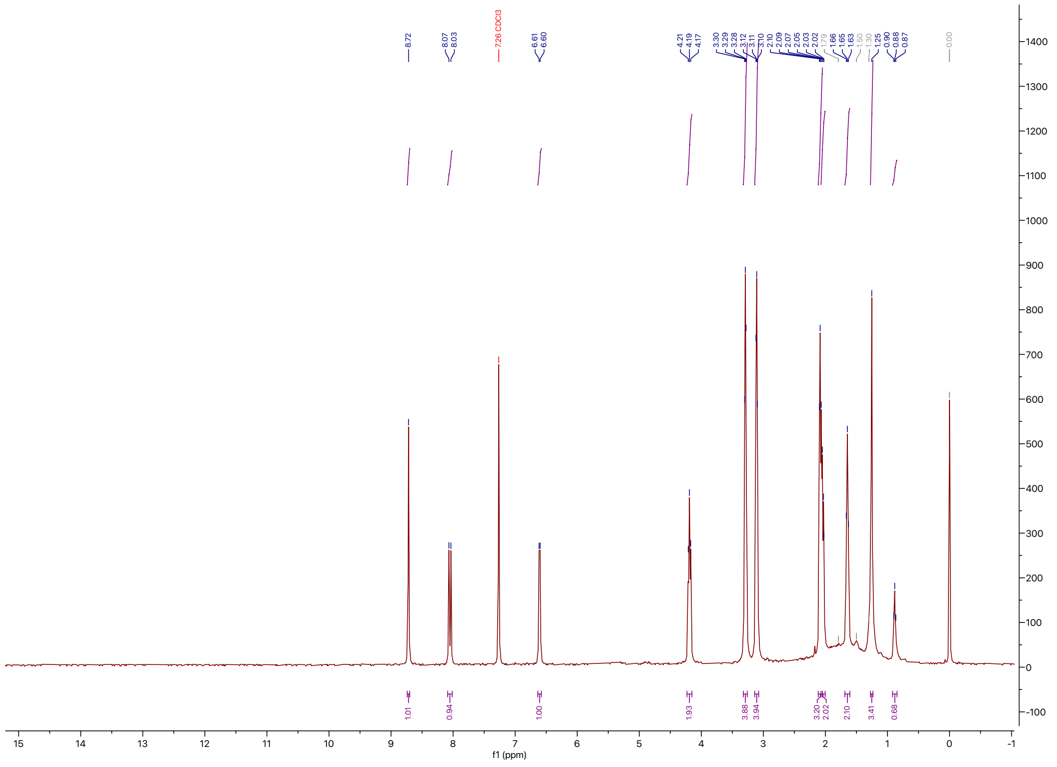

$^1$H NMR (400 MHz, $CDCl_3$) at 298 K of **2**.

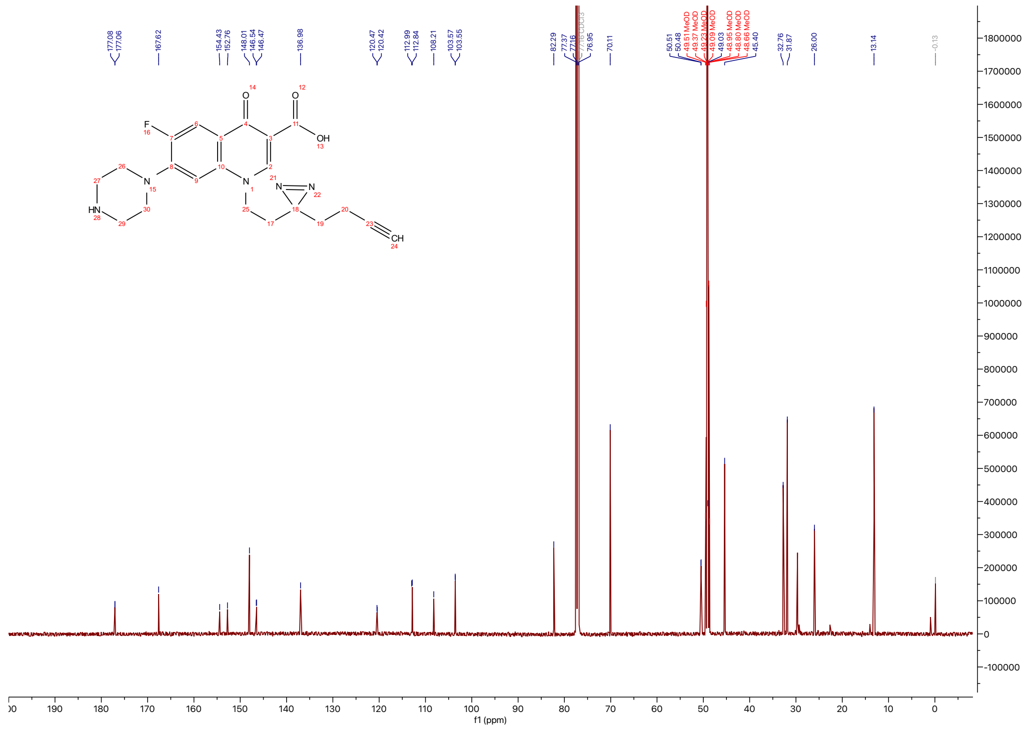

$^{13}C$ NMR (151 MHz, $CDCl_3$:MeOD) at 298 K of **2**.

***6-fluoro-4-oxo-7-(4-(prop-2-yn-1-yl)piperazin-1-yl)-1-(4-(3-(trifluoromethyl)-3H-diazirin-3-yl)benzyl)-1,4-dihydroquinoline-3-carboxylic acid*** (3)

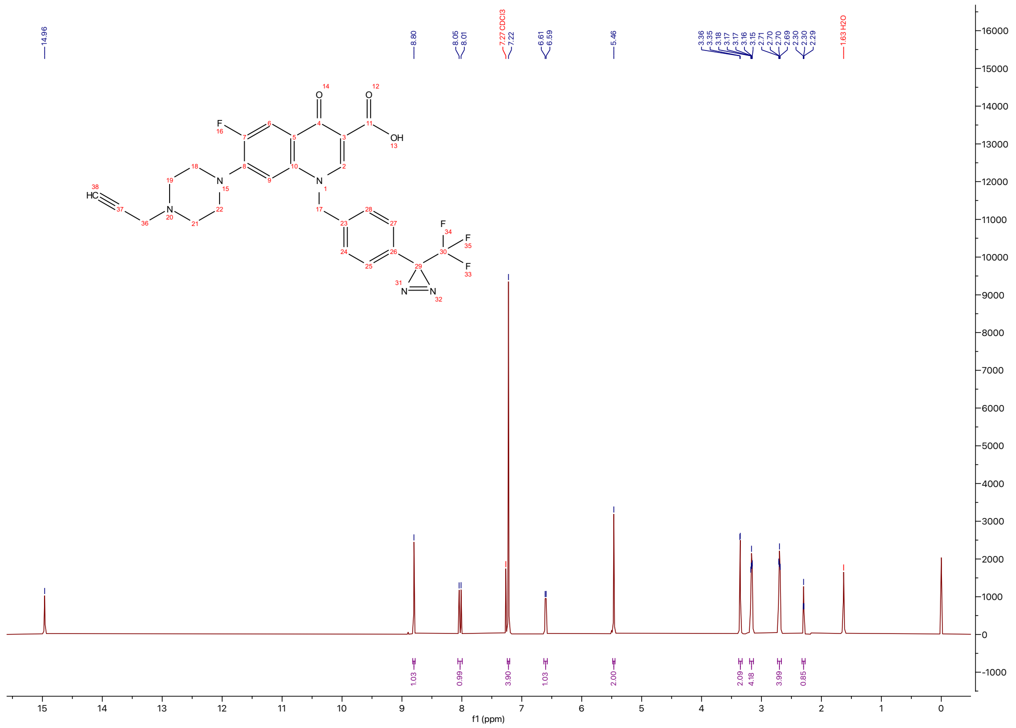

$^{1}H$ NMR (400 MHz, $CDCl_3$) at 298 K of **3**.

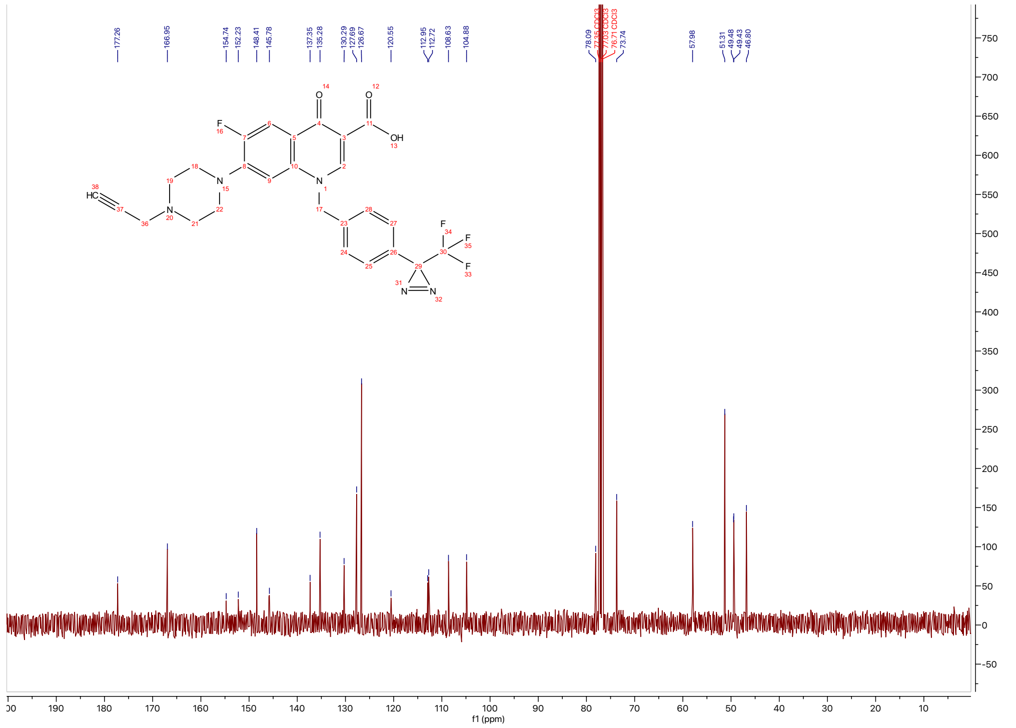

$^{13}C$ NMR (151 MHz, $CDCl_3$) at 298 K of **3**.

***6-fluoro-1-(2-methylbut-3-yn-2-yl)-4-oxo-7-(4-(4-(3-(trifluoromethyl)-3H-diazirin-3-yl)benzyl)piperazin-1-yl)-1,4-dihydroquinoline-3-carboxylic acid*** (**4**)

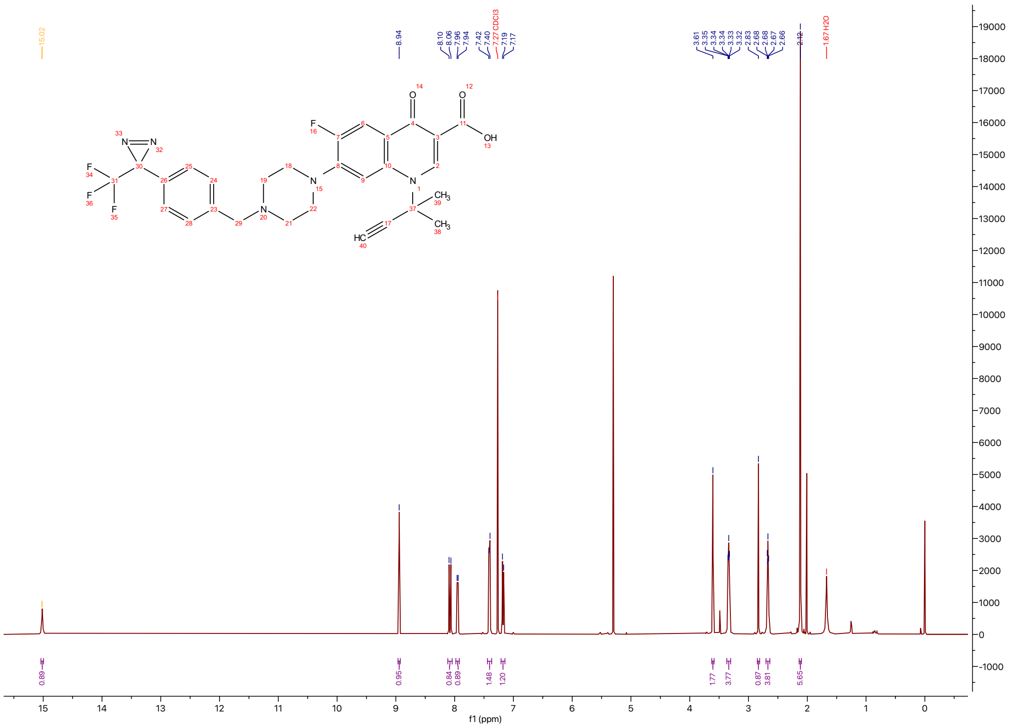

$^{1}$H NMR (400 MHz, $CDCl_3$) at 298 K of **4**.

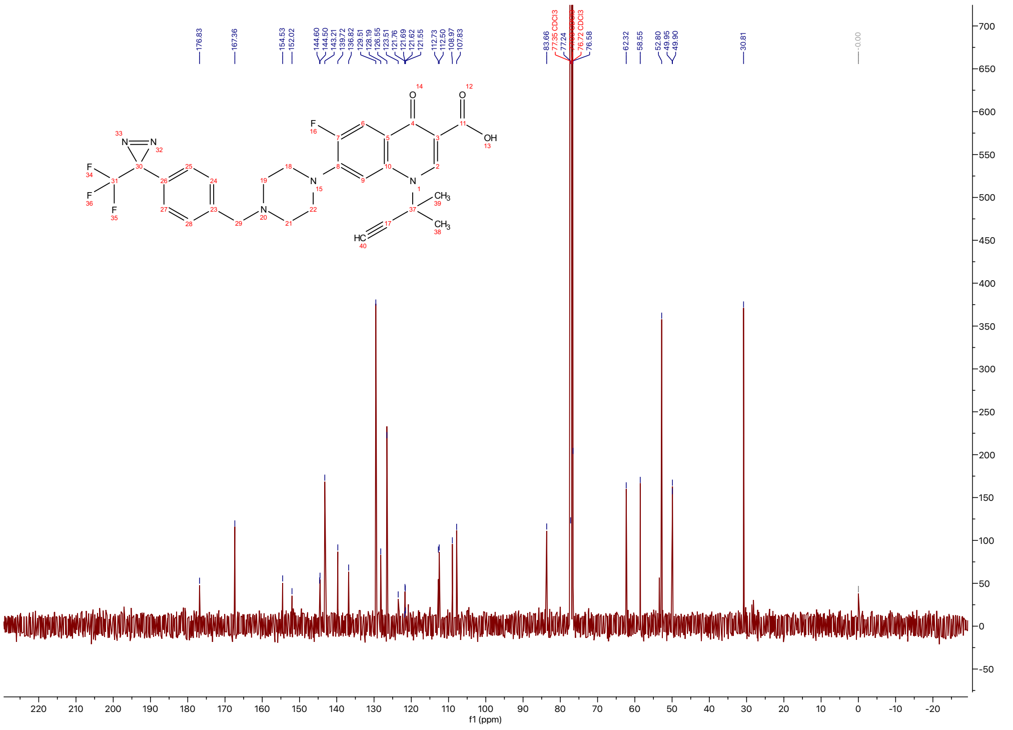

$^{13}$C NMR (151 MHz, CDCl3) at 298 K of **4**.

***7-(4-(2-(3-(but-3-yn-1-yl)-3H-diazirin-3-yl)ethyl)piperazin-1-yl)-1-cyclopropyl-6-fluoro-4-oxo-1,4-dihydroquinoline-3-carboxylic acid*** (**5**)

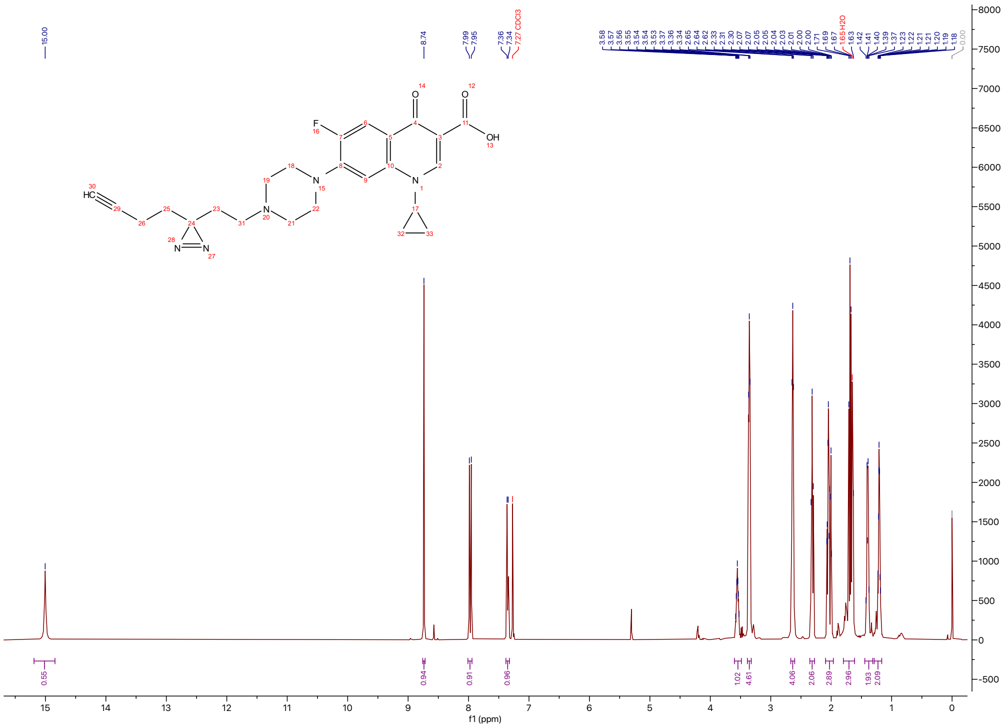

$^{1}$H NMR (400 MHz, $CDCl_3$) at 298 K of **5**.

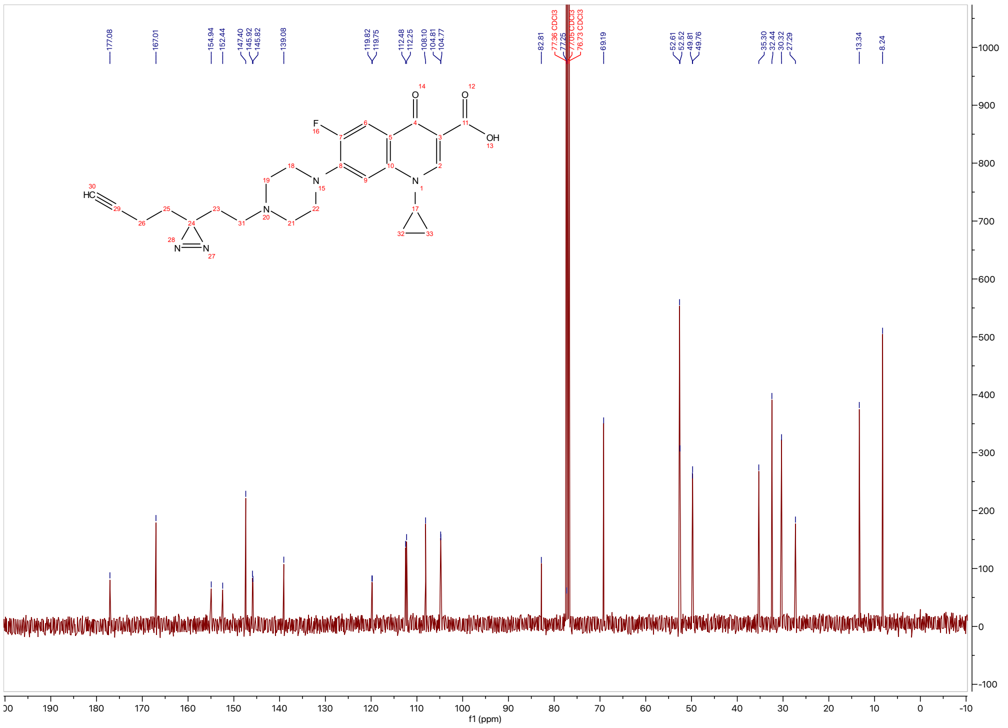

$^{13}$C NMR (151 MHz, $CDCl_3$) at 298 K of **5**.

***1-Cyclopropyl-7-(dimethylamino)-6-fluoro-4-oxo-1,4-dihydroquinoline-3-carboxylic acid (6)***

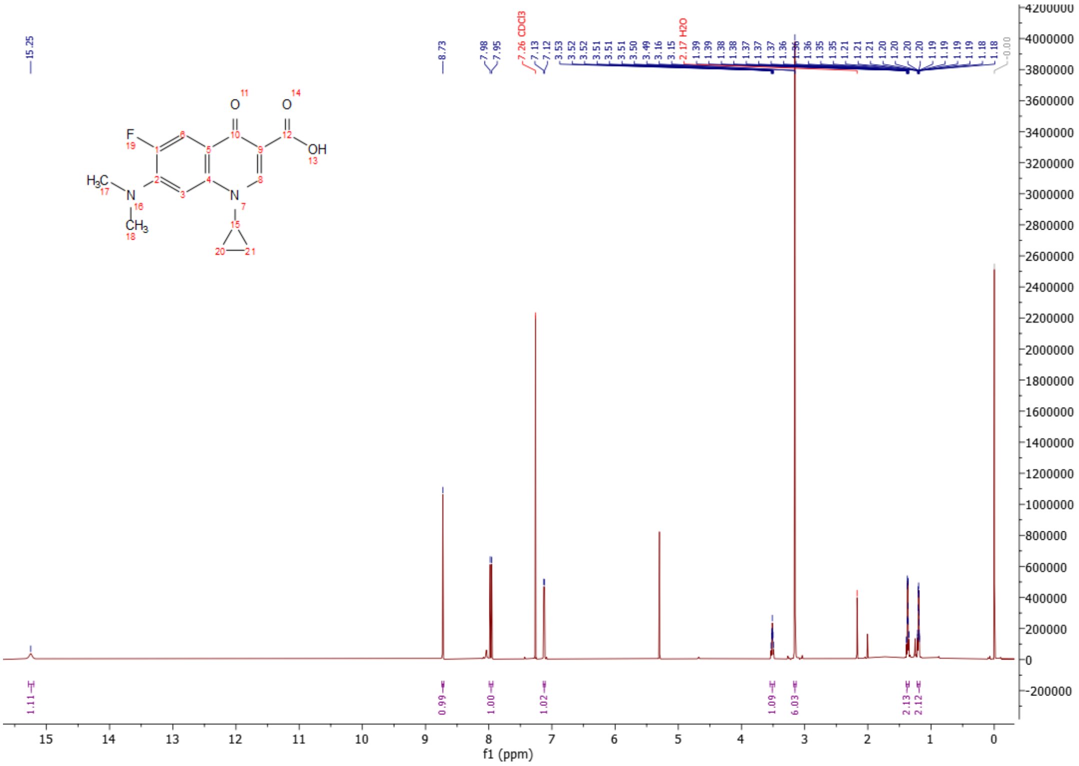

$^{1}$H NMR (600 MHz, $CDCl_3$) at 298 K of **6**.

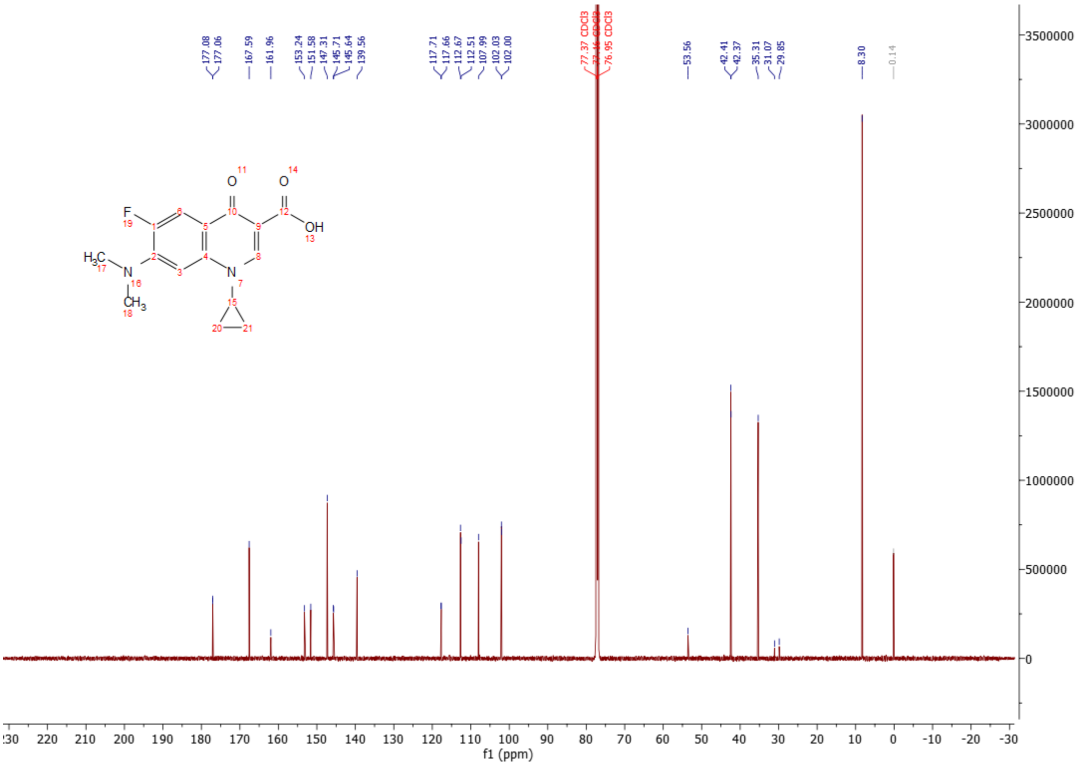

$^{13}$C NMR (151 MHz, $CDCl_3$) at 298 K of **6**.

**HPLC Traces of Final Compounds**

***1-(2-(3-(but-3-yn-1-yl)-3H-diazirin-3-yl)ethyl)-6-fluoro-4-oxo-7-(piperazin-1-yl)-1,4-dihydroquinoline-3-carboxylic acid*** (**2**)

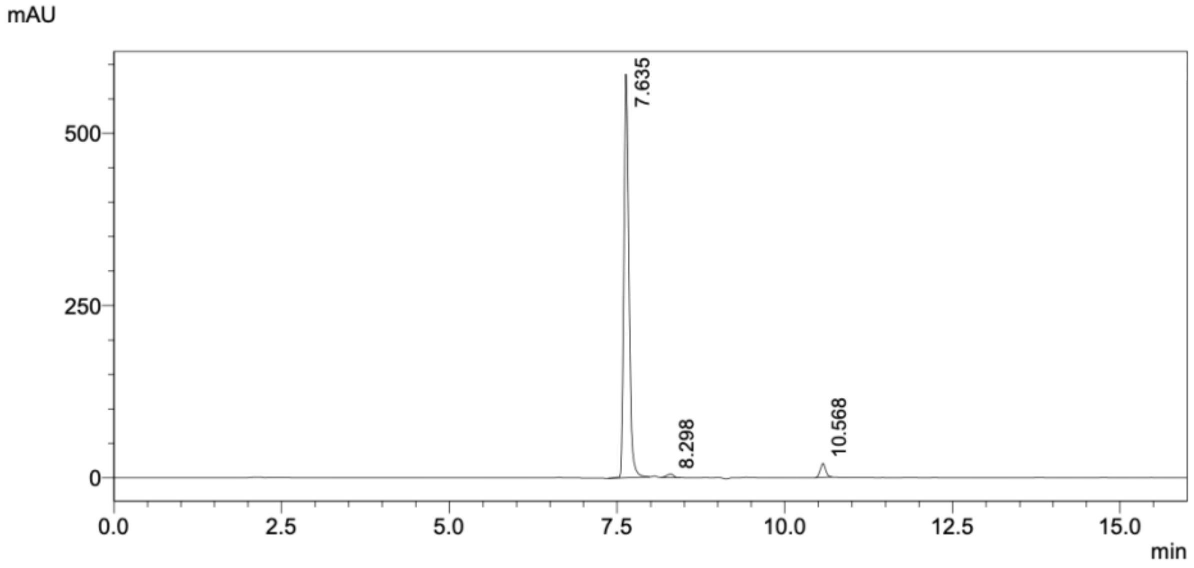

Peak Table (280 nm)

| Peak# | Ret. Time | Area | Height | Conc. | Unit | Mark | Name |
|---|---|---|---|---|---|---|---|
| 1 | 7.635 | 3057348 | 585662 | 95.425 | | M | |
| 2 | 8.298 | 34945 | 5155 | 1.091 | | M | |
| 3 | 10.568 | 111623 | 20455 | 3.484 | | M | |
| Total | | 3203915 | 611272 | | | | |

***6-fluoro-4-oxo-7-(4-(prop-2-yn-1-yl)piperazin-1-yl)-1-(4-(3-(trifluoromethyl)-3H-diazirin-3-yl)benzyl)-1,4-dihydroquinoline-3-carboxylic acid*** (**3**)

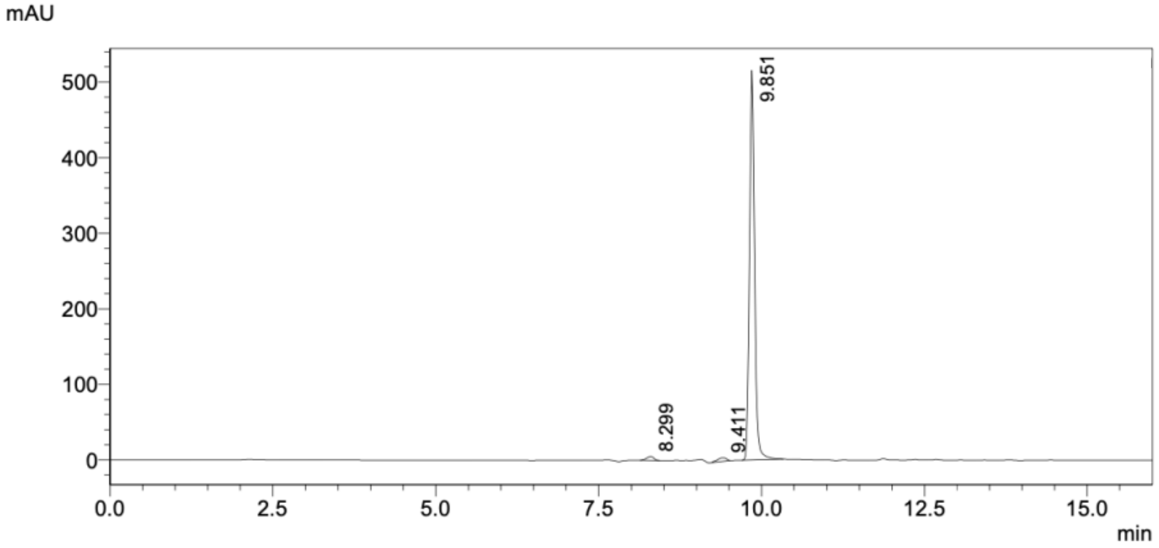

Peak Table (280 nm)

| Peak# | Ret. Time | Area | Height | Conc. | Unit | Mark | Name |
|---|---|---|---|---|---|---|---|
| 1 | 8.299 | 40003 | 5243 | 1.377 | | M | |
| 2 | 9.411 | 47393 | 4671 | 1.631 | | M | |
| 3 | 9.851 | 2817575 | 515421 | 96.992 | | M | |
| Total | | 2904970 | 525334 | | | | |

***6-fluoro-1-(2-methylbut-3-yn-2-yl)-4-oxo-7-(4-(4-(3-(trifluoromethyl)-3H-diazirin-3-yl)benzyl)piperazin-1-yl)-1,4-dihydroquinoline-3-carboxylic acid*** (**4**)

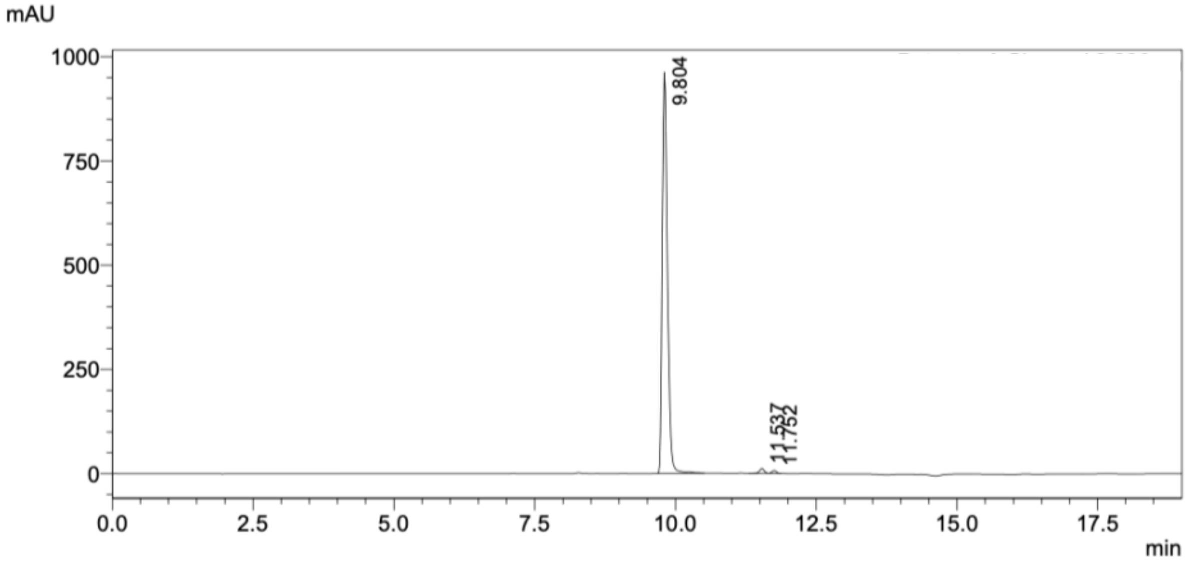

Peak Table (280 nm)

| Peak# | Ret. Time | Area | Height | Conc. | Unit | Mark | Name |
|---|---|---|---|---|---|---|---|
| 1 | 9.804 | 6176521 | 962102 | 98.344 | | M | |
| 2 | 11.537 | 67252 | 11505 | 1.071 | | M | |
| 3 | 11.752 | 36771 | 6891 | 0.585 | | M | |
| Total | | 6280544 | 980498 | | | | |

***7-(4-(2-(3-(but-3-yn-1-yl)-3H-diazirin-3-yl)ethyl)piperazin-1-yl)-1-cyclopropyl-6-fluoro-4-oxo-1,4-dihydroquinoline-3-carboxylic acid*** (**5**)

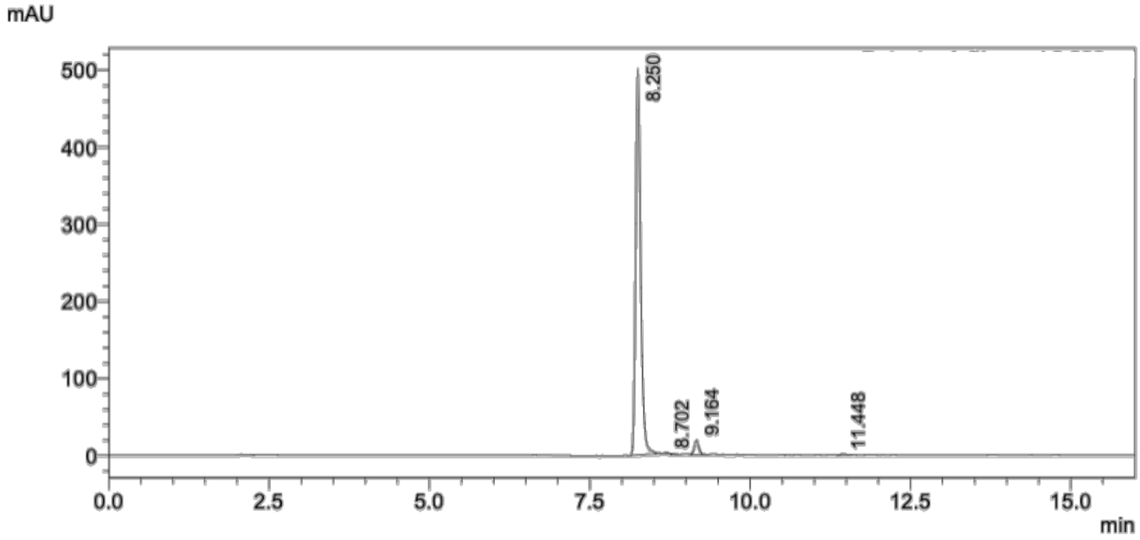

Peak Table (280 nm)

| Peak# | Ret. Time | Area | Height | Conc. | Unit | Mark | Name |
|---|---|---|---|---|---|---|---|
| 1 | 8.250 | 2792888 | 500575 | 96.451 | | M | |
| 2 | 8.702 | 5080 | 1268 | 0.175 | | M | |
| 3 | 9.164 | 90014 | 18305 | 3.109 | | M | |
| 4 | 11.448 | 7681 | 1330 | 0.265 | | M | |
| Total | | 2895663 | 521478 | | | | |

***1-Cyclopropyl-7-(dimethylamino)-6-fluoro-4-oxo-1,4-dihydroquinoline-3-carboxylic acid (6)***

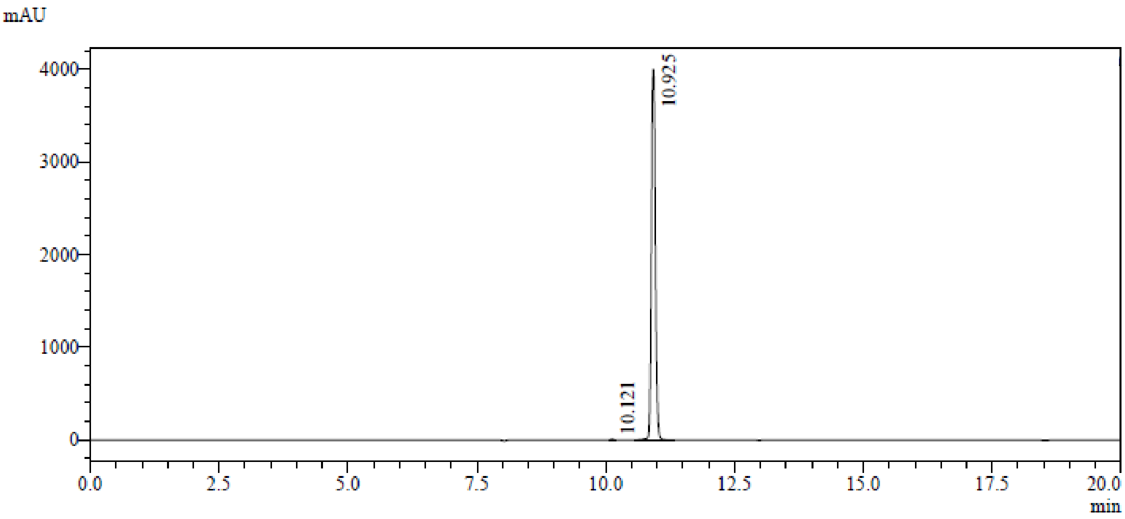

Peak Table (280 nm)

| Peak# | Ret. Time | Area | Height | Conc. | Unit | Mark | Name |
|---|---|---|---|---|---|---|---|
| 1 | 10.121 | 28050 | 7179 | 0.126 | | M | |
| 2 | 10.925 | 22153549 | 3995978 | 99.874 | | M | |
| Total | | 22181599 | 4003157 | | | | |

### References

[1] K. M. Orritt, L. Feng, J. F. Newell, J. N. Sutton, S. Grossman, T. Germe, L. R. Abbott, H. L. Jackson, B. K. L. Bury, A. Maxwell, M. J. McPhillie, C. W. G. Fishwick, *RSC Med Chem* **2022**, *13*, 831-839.

[2] Y. Y. Cheng, Z. Zhou, J. M. Papadopoulos, J. D. Zuke, T. G. Falbel, K. Anantharaman, B. M. Burton, O. S. Venturelli, *Mol. Syst. Biol.* **2023**, *19*, e11406.

[3] T. Baba, T. Ara, M. Hasegawa, Y. Takai, Y. Okumura, M. Baba, K. A. Datsenko, M. Tomita, B. L. Wanner, H. Mori, *Mol. Syst. Biol.* **2006**, *2*, 2006 0008.

[4] A. T. Kong, F. V. Leprevost, D. M. Avtonomov, D. Mellacheruvu, A. I. Nesvizhskii, *Nat. Methods* **2017**, *14*, 513-520.

[5] M. E. Ritchie, B. Phipson, D. Wu, Y. Hu, C. W. Law, W. Shi, G. K. Smyth, *Nucleic Acids Res.* **2015**, *43*, e47.

[6] J. D. Bradbury, T. Hodgkinson, A. M. Thomas, O. Tanwar, G. La Monica, V. V. Rogga, L. J. Mackay, E. K. Taylor, K. Gilbert, Y. Zhu, A. Y. Sefton, A. M. Edwards, C. J. Gray-Hammerton, G. R. Smith, P. M. Roberts, T. R. Walsh, T. Lanyon-Hogg, *Chem Sci* **2024**, *15*, 9620-9629.
